# Conventional NK cells and tissue-resident ILC1s join forces to control liver metastasis

**DOI:** 10.1101/2020.07.17.206433

**Authors:** Laura Ducimetière, Giulia Lucchiari, Gioana Litscher, Marc Nater, Laura Heeb, Nicolás Gonzalo Nuñez, Laura Wyss, Dominik Burri, Marijne Vermeer, Julia Gschwend, Andreas E. Moor, Burkhard Becher, Maries van den Broek, Sonia Tugues

## Abstract

The liver is a major metastatic target organ, and little is known about the role of immunity in controlling hepatic metastases. Here, we discovered that the concerted and non-redundant action of two innate lymphocyte subpopulations, conventional NK cells (cNKs) and tissue-resident type I Innate Lymphoid Cells (trILC1s), is essential for anti-metastatic defense. Using different preclinical models for liver metastasis, we found that trILC1 control metastatic seeding, whereas cNKs restrain outgrowth. The antimetastatic activity of cNKs is regulated in a tumor type-specific fashion. Thereby, individual cancer cell lines orchestrate the emergence of cNK subsets with unique phenotypic and functional traits. Understanding cancer-cell- as well as innate-cell-intrinsic factors will allow the exploitation of hepatic innate cells for development of novel cancer therapies.

**Significance:** Innate lymphoid cells hold great promise for the treatment of metastases. Development of effective therapies based on these versatile immune cells, however, is hampered by our limited knowledge of their behavior in the metastatic niche. Here, we describe that defense against liver metastasis requires the collaboration between two innate lymphocyte subsets, conventional NK cells (cNKs) and tissue-resident type I innate lymphoid cells (trILC1s). We show that different cancers generate their own particular metastatic niche inducing specific changes in cNKs and trILC1s. Further, we uncover specific cNK subsets that can be manipulated to improve their anti-metastatic potential. Our work contributes to understanding how cancer-specific factors and hepatic innate lymphocytes exert mutual influence and how this can be exploited for therapeutic purposes.

**Highlights:**

- cNKs and trILC1s collaborate to control hepatic metastasis
- trILC1s restrict seeding and cNKs control outgrowth of cancer cells in the liver
- Individual cancer cell lines orchestrate a distinct metastatic niche
- The metastatic niche dictates the phenotype and function of cNKs

## INTRODUCTION

Hepatic metastases are a major cause of cancer-related death in disparate cancer types (Williamson et al., 2019). The healthy liver contains a large population of innate lymphocytes including conventional natural killer cells (cNKs) and tissue-resident type I innate lymphoid cells (trILC1s). Although hepatic cNKs and trILC1s share some features, they differ in phenotype, ontogeny and function (Peng et al., 2013; Sojka et al., 2014; Takeda et al., 2005). In steady state, both cNKs and trILC1s are defined as Lin^-^ CD45^+^NK1.1^+^NKp46^+^ lymphocytes, with cNKs being CD49a^-^CD49b^+^ and trILC1s CD49a^+^CD49b^-^. Whereas cNKs rely on the transcription factors Eomes (Gordon et al., 2012) and Nfil3 (Kamizono et al., 2009) for their development, trILC1s rather depend on T-bet (Daussy et al., 2014), Hobit (Mackay et al., 2016) and the aryl hydrocarbon receptor (AhR) (Zhang et al., 2016). Further, cNKs express higher amounts of markers associated with maturation (CD11b, CD43, KLRG-1) compared to trILC1s, which rather feature the cytotoxic molecule TNF-related apoptosis-inducing ligand (TRAIL) and markers for tissue-residency (e.g. CD69) (Fuchs, 2016). Upon activation, both cNKs and trILC1s produce IFN-γ and are cytotoxic (Ishiyama et al., 2006; Takeda et al., 2005). However, trILC1s express more inhibitory receptors (Peng et al., 2013; Sojka et al., 2014), suggesting they perform immune-regulatory functions. Nevertheless, the main roles ascribed to trILC1s so far include immune responses against haptens and early protection against viral antigens and acute liver injury (Nabekura et al., 2020; Peng et al., 2013; Weizman et al., 2017).

The protective role of NK cells against spontaneous and experimental metastasis to the lungs is well established (Beffinger et al., 2018; Chiossone et al., 2018; Guillerey and Smyth, 2016; López-Soto et al., 2017; Ohs et al., 2016). At this particular site, cNKs progressively lose their protective capacities, which limits their anti-metastatic effect (Carrega et al., 2008; Ohs et al., 2017; Platonova et al., 2011). Whether innate lymphocytes control hepatic metastases and the individual contributions of cNKs and trILC1s is largely unknown.

Understanding of innate immune defense against liver metastasis is pivotal to exploiting innate lymphocytes for novel cancer therapies. Here we sought to unravel the mechanism that governs metastatic control in this immunologically unique organ, and discovered that both cNKs and trILC1s controlled the metastatic burden in the liver, but seemed to have non-redundant roles. Whereas trILC1s emerged as important regulators of metastatic seeding, the cNK cell subset had a greater potential to clear cancer cells throughout metastatic progression. The function and phenotype of cNKs were regulated in a cancer-type specific fashion. Hence, cNKs responded differently to factors produced in the metastatic niche leading to the emergence of unique cNK cell subsets.

## RESULTS

### Control of hepatic metastases depends on NKp46^+^ innate cells

Despite the well-accepted role of NK cells in the control of tumor progression, their influence on hepatic metastases is still poorly understood. We used two different models for liver metastasis to address this question: MC38 colon carcinoma cells and Lewis Lung Carcinoma cells (LLC). We selected these two models based on their intrinsic difference in the activation of innate immune pathways. Whereas MC38 cells produce cyclic GMP-AMP (cGAMP) resulting in the local production of type I interferon (IFN) and consequent activation of NK and CD8^+^ T cells, LLC do not produce cGAMP and consequently, do not activate downstream pathways (Marcus et al., 2018; Schadt et al., 2019). We induced liver metastasis from MC38 or LLC carcinoma cells and depleted NKp46^+^ (encoded by *Ncr1*) subsets at different time points by administering diphtheria toxin (DTX) to *Ncr1*^iCre^*R26R*^iDTR^ mice (Figures 1A, 1B and Supplementary Figure S1A). The metastatic load increased dramatically in both models when NKp46^+^ cells were depleted 48 h before or 24 h after tumor cell injection (Figures 1C and 1D). However, their depletion 7 days after tumor injection resulted in larger liver metastases from MC38 but not LLC cells (Figures 1C and 1D). Thus, NKp46^+^ cells control seeding of MC38 and LLC cells but only the outgrowth of MC38 cells, suggesting that they become ineffective against progressing LLC metastases. In this experimental set up, CD8^+^ T cells are dispensable for restricting liver metastases (Supplementary Figure S1B).

**Figure 1:**
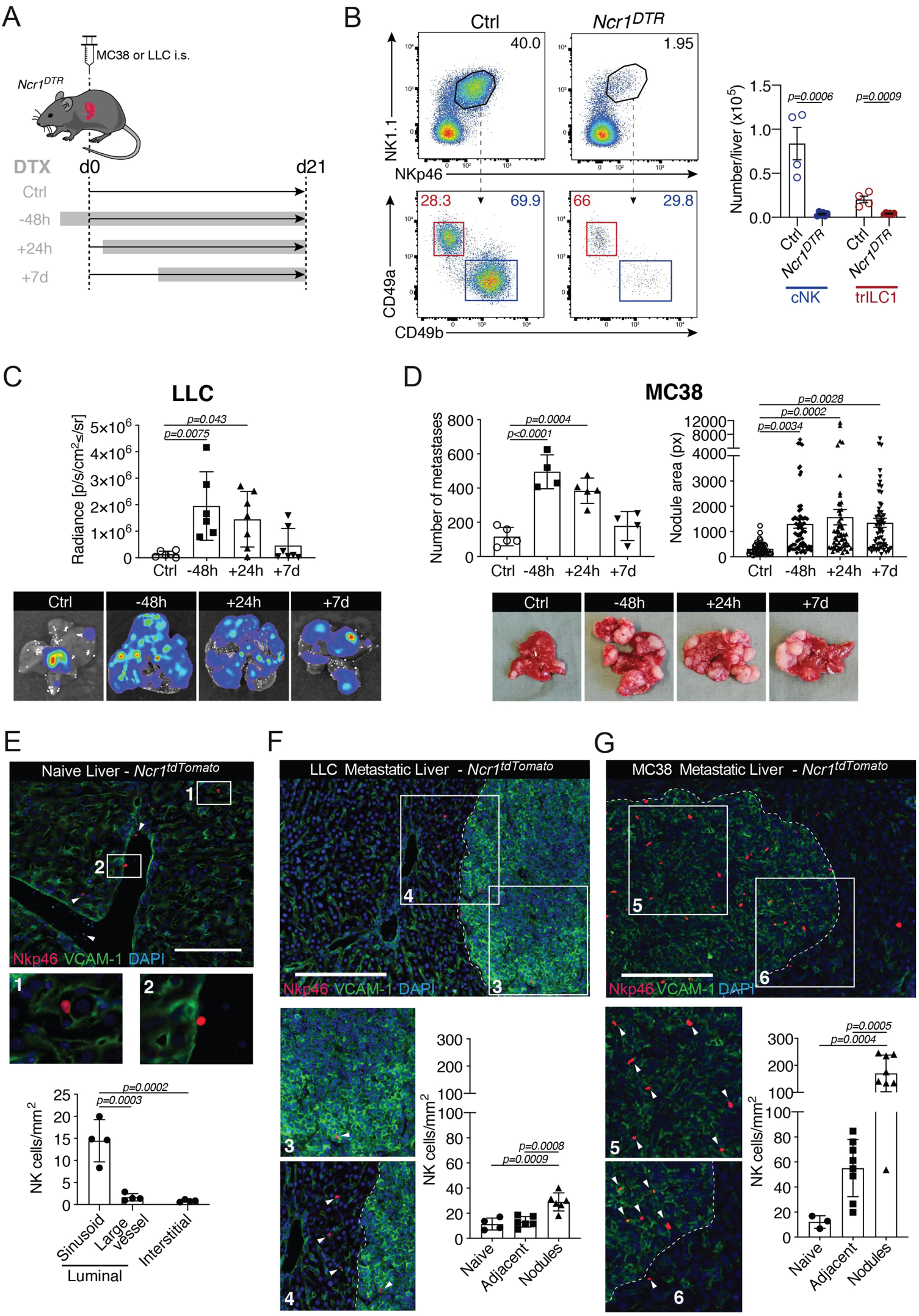
Control of hepatic metastases depends on NK cells. **(A)** Experimental schedule for NK depletion. NK cells were depleted by injection of 250 ng diphtheria toxin intraperitoneally 48 hours before, 24 hours after or 7 days after tumor injection. Depletion was maintained until the endpoint. *Ncr1*^*DTR*^ = *Ncr1*^*iCre/wt*^.*R26R*^*iDTR/wt*^; Ctrl = undepleted control mice (*Ncr1*^*iCre/wt*^.*R26R*^*wt/wt*^, group injected at time point -48h); i.s. = intra-splenic; DTX = diphtheria toxin. **(B)** Representative dot plots (left) and quantification (right) of cNKs and trILC1s in Ctrl and *Ncr1*^*DTR*^ livers. Samples were pre-gated on single live CD45^+^lineage^-^ cells and subsequently gated on NK1.1^+^NKp46^+^ cells. cNK = conventional NK cells, CD49a^-^ CD49b^+^; trILC1 = tissue-resident ILC1s, CD49a^+^CD49b^-^. **(C)** Upper panel: Quantification of LLC liver nodules by *ex vivo* IVIS imaging at the endpoint. Lower panels: Representative IVIS images of metastatic livers from each group at the endpoint. The bar represents the mean ± SD, symbols represent livers from individual mice. One-way analysis of variance (ANOVA), with Tukey’s multiple comparisons test. The experiment was performed twice with similar results. **(D)** Upper panel: Quantification of MC38 macroscopic liver nodules and their area at the endpoint. Lower panels: Representative images of metastatic livers from each group at the endpoint. The bar represents the mean ± SD, symbols represent livers from individual mice. One-way analysis of variance (ANOVA), with Tukey’s multiple comparisons test. The experiment was performed twice with similar results. **(E-G)** Representative immunofluorescence image and quantification of NK cell vascular localization. *Ncr1*^*tdTomato*^ = *Ncr1*^*Cre/wt*^.*R26R*^*Ai14/wt*^. Three livers per experimental condition were analyzed, each symbol represents an individual section. **(E)** Naïve liver. (**F)** LLC-metastatic liver. **(G)** MC38-metastatic liver. The scale bar corresponds to 50 µm (E) or 100 µm (F, G).

Because NKp46^+^ cells lose control over LLC metastases after some time, we investigated the location of NK cells within the hepatic MC38 and LLC metastatic niche. For this purpose, we used *Ncr1*^iCre^*Tdtomato*^fl/wt^ mice, in which NKp46^+^ cells are irreversibly marked with TdTomato (Supplementary Figure S1C), and visualized tumor cells and the sinusoidal endothelium with an antibody against VCAM-1. In healthy livers, NKp46^+^ cells were mostly located along the luminal aspect of the sinusoids (Figure 1E). NK cells were abundant in MC38 nodules, whereas they were sparse in LLC nodules (Figures 1F and 1G). Thus, failure to infiltrate LLC nodules may explain why NKp46^+^ cells are relatively ineffective in controlling disseminated LLC cells in the liver.

Together, these findings highlight the crucial contribution of cNKs and trILC1s to control of hepatic metastasis and reveal important differences linked to cancer-cell-intrinsic traits.

### Division of labor between hepatic cNKs and trILC1s in the control of metastasis

The liver contains a high number of cNKs and trILC1s. The relative contribution of these two innate cell types to metastatic surveillance is not known. Generally, CD49a^-^ CD49b^+^ cNKs are phenotypically more mature and differentiated, while CD49a^+^CD49b^-^ trILC1s have a combined signature of activation and immaturity (Peng et al., 2013). To determine the individual contribution of cNKs and trILC1s to the control of hepatic metastases we used two genetically-modified mouse lines that lack one of the two innate subsets: *Ncr1*^iCre^*Eomes*^fl/fl^ mice (Gordon et al., 2012; Weizman et al., 2017) severely reduced in cNKs but not trILC1s (Figure 2A) and Hobit-deficient (*Hobit*^-/-^) mice (Mackay et al., 2016), selectively lacking trILC1s but not cNKs (Figure 2B). We found that both cNK and trILC1s restricted the progression of MC38- and LLC-derived liver metastasis, as shown by the higher metastatic load in *Ncr1*^iCre^*Eomes*^fl/fl^ and *Hobit*^-/-^ livers in comparison to their respective littermate controls (Figures 2C and 2D). We corroborated the anti-metastatic properties of cNKs using *Nfil3*^-/-^ mice, which are also devoid of cNKs (Gascoyne et al., 2009) (Supplementary Figures S1D and S1E). The difference between LLC and MC38 metastases concerning the time window of NK-dependent control led us to investigate the temporal requirements for cNKs and trILC1s to exert their anti-metastatic activity. Therefore, we first crossed mice missing cNKs (*Ncr1*^iCre^*Eomes*^fl/fl^) to *R26R*^iDTR^ mice and additionally eliminated trILC1s via DTX administration at different time points relative to tumor cell injection as shown in Figure 1A. Depletion of trILC1s 48 h prior to tumor cell injection – but not thereafter – resulted in an increased load of MC38- or LLC-derived metastases (Figures 3A and 3B), indicating that trILC1s mainly control metastatic seeding. In agreement, trILC1s poorly infiltrate MC38-derived metastatic nodules, as shown when using *Ncr1*^iCre^*Eomes*^fl/fl^ *TdTomato*^fl/wt^ mice (Figure 3C). To study the anti-metastatic potential of cNKs, we performed a similar time-course experiment in trILC1-deficient *Hobit*^-/-^ mice and depleted cNKs using anti-asialo-GM1 antibodies (as shown in Figure 1A). cNKs controlled the early stages of both LLC- and MC38-derived metastasis (Figure 3D). In contrast, however, cNKs were ineffective against late-stage LLC nodules but still restricted (Figure 3E) and infiltrated late-stage MC38 nodules (Figure 3F). These findings agree with what we observed using *Nkp46*^iCre^*R26R*^iDTR^ mice, and suggest that particularly the long-term anti-metastatic effect of cNK is influenced by cancer-cell-intrinsic features and/or the tumor microenvironment.

Collectively, these results point towards an important role of trILC1s in limiting metastatic seeding. cNKs, however, are more prone to infiltrate metastatic nodules and provide an enduring anti-metastatic activity.

**Figure 2:**
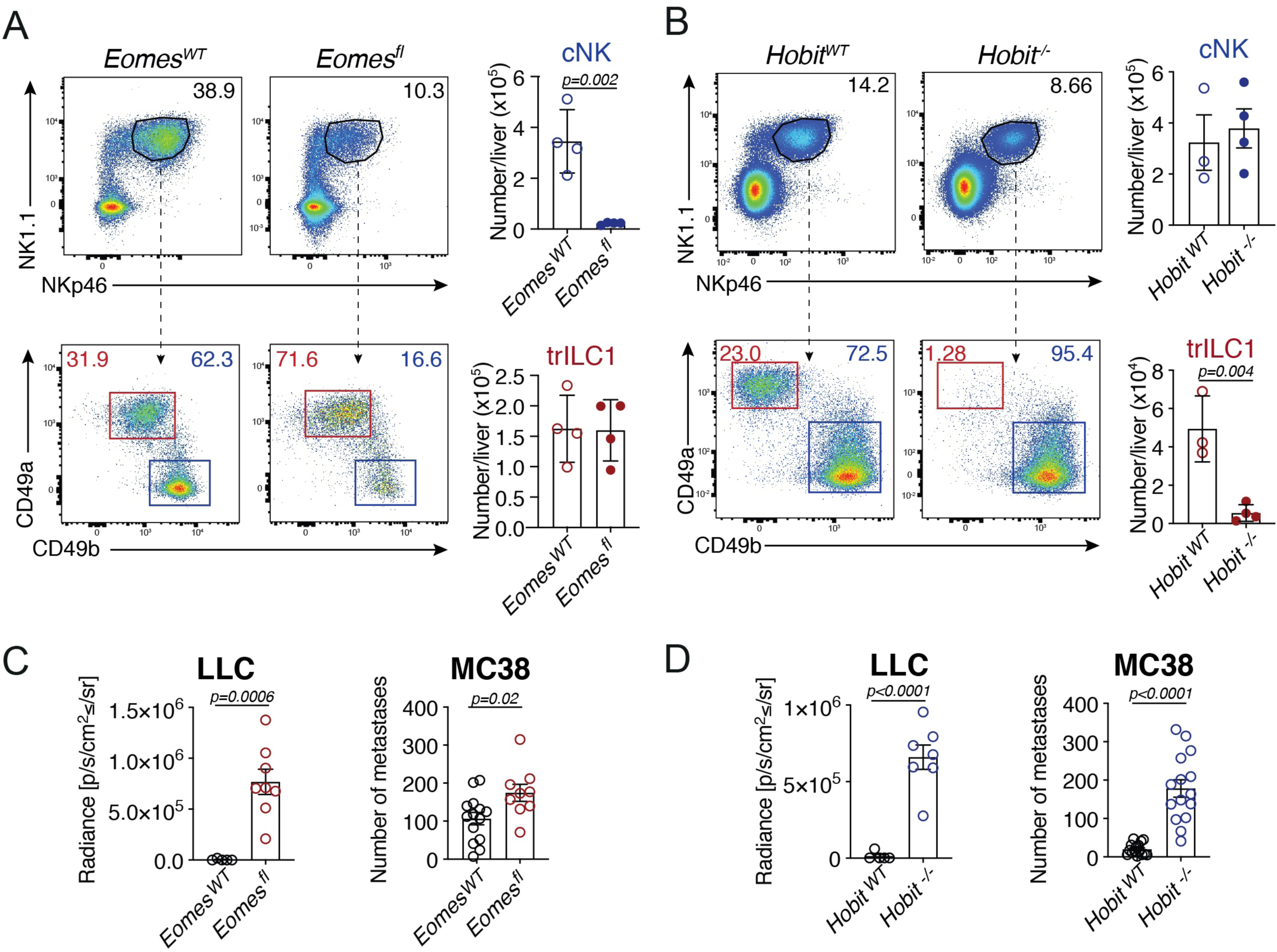
Division of labor between hepatic cNKs and trILC1s in the control of metastasis. **(A, B)** Representative dot plots and quantification of cNKs and trILC1s. Samples were pre-gated on single live CD45^+^lineage^-^ cells and subsequently gated on NK1.1^+^NKp46^+^ cells. cNK = conventional NK cells, CD49a^-^CD49b^+^ ; trILC1 = tissue-resident ILC1s, CD49a^+^CD49b^-^. **(A)** *Eomes*^*WT*^ and *Eomes*^*fl*^ livers. *Eomes*^*WT*^ = *Nkp46*^*iCre/wt*^.*Eomes*^*wt/wt*^; *Eomes*^*fl*^ = *Nkp46*^*iCre/wt*^.*Eomes*^*fl/fl*^. **(B)** *Hobit*^*WT*^ and *Hobit*^-/-^ livers. (**C, D)** Quantification of the metastatic burden from LLC and MC38 cells in **(C)** *Eomes*^*WT*^ and *Eomes*^*fl*^ mice and **(D)** *Hobit*^*WT*^ and *Hobit*^*-/-*^ mice. The bar represents the mean ± SD, symbols represent livers from individual mice. Student’s unpaired t test. LLC: The experiment was performed twice with similar results. MC38: Pooled data from 4 experiments.

**Figure 3:**
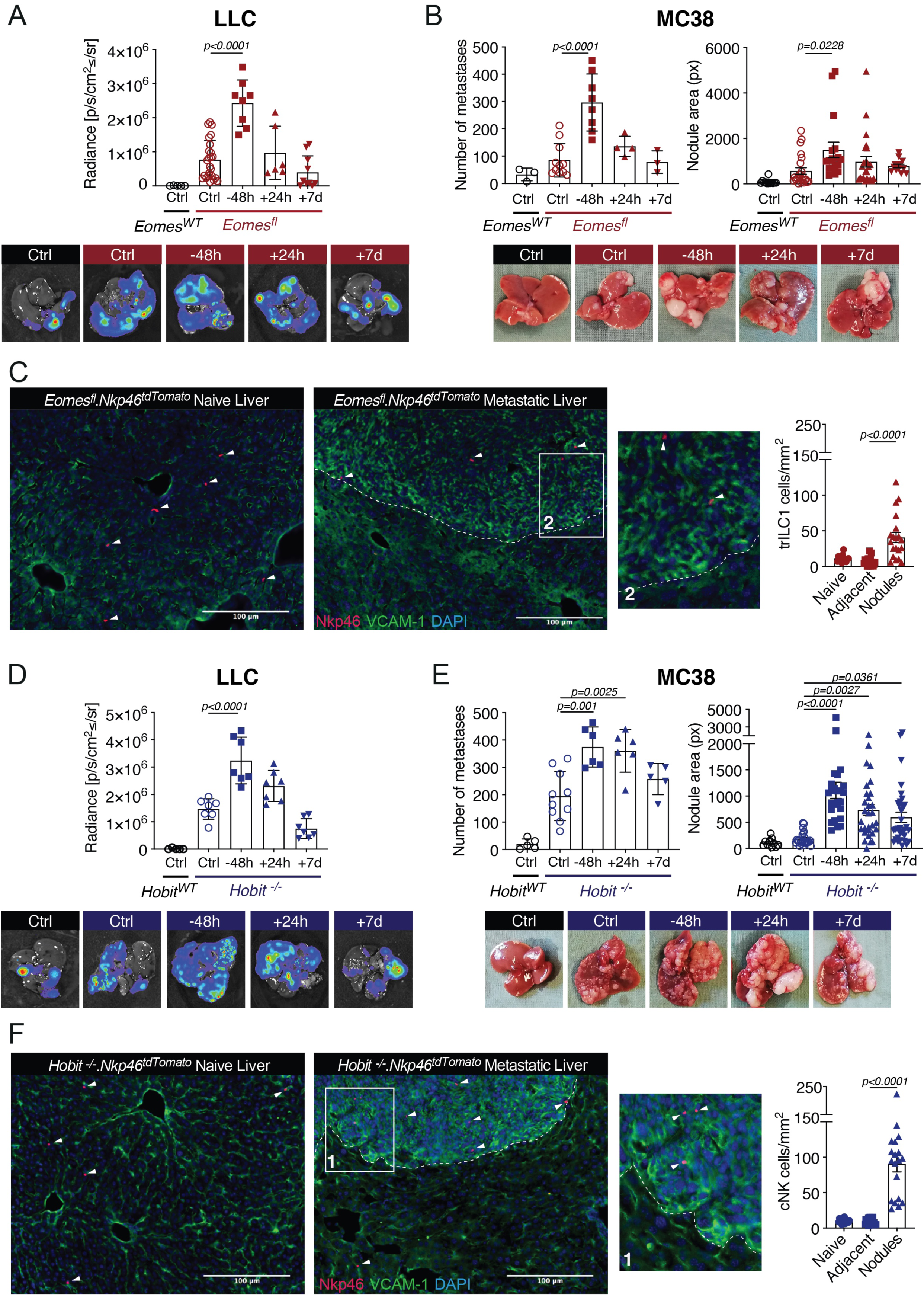
cNKs but not trILC1s control advanced metastatic disease. **(A, B)** cNKs were depleted by injection of 250 ng diphtheria toxin intraperitoneally 48 hours before, 24 hours after or 7 days after tumor injection. Depletion was maintained until the endpoint (day 18). *Eomes*^*WT*^ Ctrl = *Ncr1*^*iCre/wt*^.*Eomes*^*wt/wt*^.*R26R*^*wt/wt*^; *Eomes*^*fl*^ Ctrl = *Ncr1*^*iCre/wt*^.*Eomes*^*fl/fl*^.*R26R*^*wt/wt*^; *Eomes*^*fl*^ NK-depleted = *Ncr1*^*iCre/wt*^.*Eomes*^*fl/fl*^.*R26R*^*iDTR/wt*^. Upper panels: Quantification of liver nodules at endpoint. Lower panels: Representative images of metastatic livers at endpoint. The bar represents the mean ± SD, symbols represent livers from individual mice. One-way analysis of variance (ANOVA), with Tukey’s multiple comparisons test. The experiment was performed twice with similar results. **(A)** LLC-liver nodules. **(B)** MC38-liver nodules. **(C)** Left three panels: Representative immunofluorescence images of a naïve and MC38-metastatic liver from *Ncr1*^*iCre/wt*^.*Eomes*^*fl/fl*^.*R26R*^*Ai14/wt*^ mice. Right panel: Localization of trILC1s. Three naïve and 4 MC38-metastatic livers were analyzed, each symbol represents a liver section. One-way analysis of variance (ANOVA), with Tukey’s multiple comparisons test. **(D, E)** Depletion of NK cells in *Hobit*^*WT*^ and *Hobit*^*-/-*^ mice using anti-Asialo-GM1. Depletion was started 48 hours before, 24 hours after or 7 days after tumor injection, and maintained until the endpoint (day 18). Upper panels: Quantification of liver nodules at endpoint. Lower panels: Representative images of metastatic livers at endpoint. The bar represents the mean ± SD, symbols represent livers from individual mice. One-way analysis of variance (ANOVA), with Tukey’s multiple comparisons test. The experiment was performed twice with similar results. **(D)** LLC-liver nodules. **(E)** MC38-liver nodules. **(F)** Left three panels: Representative immunofluorescence images of a naïve and MC38-metastatic liver from *Ncr1*^*iCre/wt*^.*Hobit*^*-/-*^.*R26R*^*Ai14/wt*^ mice. Right panel: Localization of cNKs. Three naïve and 3 MC38-metastatic livers were analyzed, each symbol represents a liver section. One-way analysis of variance (ANOVA), with Tukey’s multiple comparisons test.

### cNKs but not trILC1s progressively lose function in hepatic metastases

The apparent correlation between the capacity of cNKs to control established metastasis and to infiltrate the latter led us to study the functionality of cNKs in advanced metastatic disease (around 3 weeks after tumor injection). For this purpose, we compared cells isolated from naïve liver, tumor-adjacent tissue and metastatic nodules. We found impaired cytotoxicity of cNKs from MC38-and LLC-metastatic livers when compared to cNKs isolated from naive livers (Figure 4A). In contrast, trILC1s from both tumor models preserved the ability to kill tumor cells even in an environment of advanced metastasis (Figure 4B). Despite the impaired cytotoxicity in both models, cNKs from MC38-metastatic livers produced high amounts of granzyme B (GrzB), both when isolated from the adjacent tissue as well as from the nodules (Figure 4C). This was not the case for cNKs from LLC-metastatic livers, which secreted even less GrzB than their naïve counterparts (Figure 4C and Supplementary Figure S2A).

**Figure 4:**
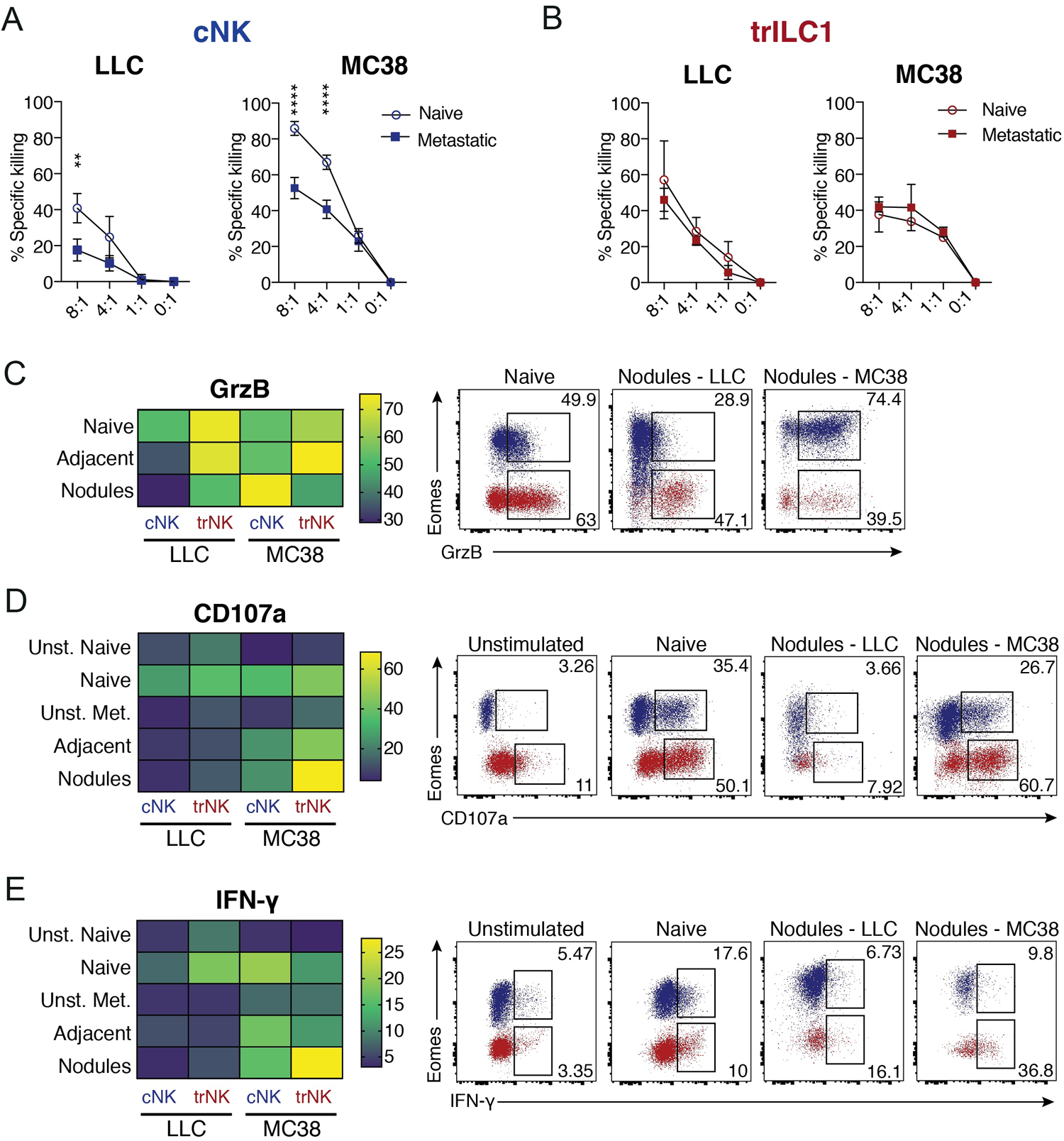
cNKs but not trILC1s progressively lose function in hepatic metastases. **(A)** cNKs and **(B)** trILC1 were sorted from naïve, LLC-metastatic or MC38-metastatic livers, and incubated with the respective cell lines at different effector:target ratios for 12 hours. Percentage specific killing was calculated as follows: [(% cell death in the presence of NK cells) — (% spontaneous cell death)] ÷ (100 — % spontaneous cell death). Results are displayed as mean of 3 replicates ± SD. Student’s unpaired t test: **p= 0.0076, ****p<0.0001. The experiment was performed twice with similar results. **(C-E)** Expression of effector molecules by NK cells in metastatic livers. Metastatic livers were manually dissected to separate the nodules from the adjacent tissue, and tissues were enzymatically processed into a single-cell suspension. Granzyme B was stained intracellularly directly *ex vivo*; CD107a and IFN-γ were stained after 4 hours of stimulation by plate-bound anti-NK1.1. Left panels: Heatmaps showing the mean percentages of positive cells. Right panels: Representative dot plots showing the expression and gating strategy for GrzB^+^, CD107a^+^ and IFN-γ^+^ cells in cNKs (blue) and trILC1s (red). Groups contained 5-6 mice. The experiment was performed twice with similar results. Flow cytometry analysis of **(C)** Granzyme B (GrzB), **(D)** CD107a and **(E)** IFN-γ expression by cNKs and trILC1s cells from naïve and metastatic livers.

Next, we investigated additional functions of cNKs and trILC1s including their capacity to degranulate (indicated by surface expression of CD107a (LAMP-1) (Betts et al., 2003) and produce IFN-γ in response to different stimuli. We found that both cNKs and trILC1s from MC38-metastatic but not from LLC-metastatic livers degranulated and produced IFN-γ in response to anti-NK1.1 stimulation (Figures 4D, 4E and Supplementary Figures S2B and S2C). We observed similar results when using PMA plus Ionomycin as a stimulus (Supplementary Figure S2D). However, the impaired IFN-γ-production of LLC-derived cNKs could be rescued by incubation with IL-15 or IL-12 plus IL-18 (Supplementary Figures S3A-S3D).

Thus, although hepatic cNKs become dysfunctional during metastasis formation, some functions are better maintained in the context of MC38 than of LLC-derived metastasis. This is in line with the inability of cNKs to control established LLC-nodules.

### Metastatic livers drive the emergence of unique cNK cell populations

To better understand the influence of the MC38- and LLC-associated hepatic metastatic niche on cNKs and trILC1s, we characterized both NK cell subsets using high-parametric single-cell cytometry. We used cell suspensions of naïve liver as well as the adjacent tissue and nodules from metastatic sites and focused on the expression of NK cell surface receptors, transcription factors and effector molecules. To better visualize phenotypical variations, we reduced the high-dimensional dataset into two dimensions using the uniform manifold approximation and projection (UMAP) algorithm (McInnes et al, J Open Source Softw. 2018). Typical UMAP plots show the normalized expression of each individual marker (Supplementary Figures S4A and S4B). We then combined this analysis to the self-organizing map (FlowSOM) metaclustering algorithm (Brummelman et al., 2019) to identify distinct populations (Figures 5A and 5H). This approach revealed two clusters of cNKs, termed cNK_1 and cNK_2, corresponding to more differentiated (CD49b^high^, CD11b^high^, Eomes^high^, Tbet^high^) and less differentiated (CD49b^int^, CD11b^low^, Eomes^int^, Tbet^int/low^) cNKs, respectively (Figures 5A-D, 5H-K and Supplementary Figure S4C). We found an expansion of cNK_2 cells in metastatic livers of both MC38 and LLC tumor bearing mice, in agreement with the increased numbers of less differentiated NK cells reported in the pulmonary metastatic niche (Ohs et al., 2017) (Figures 5B, 5C, 5I and 5J). Besides these expected phenotypes, we found unique NK cell subsets in metastatic livers that, however, differed in MC38 and LLC metastases, suggesting tumor-driven education. In LLC-metastatic livers, a cNK subset emerged that is characterized by a very low expression of CD49b, CD49a, KLRG-1, CD11b, Eomes, T-bet, GrzB and Ki67, but high expression of CD69, Thy1.2 and CD27. We refer to this subset as CD49a^-^Eomes^-^ cNKs, which are reminiscent of very immature NK cells (Figures 5B-5F and Supplementary Figure S4C). CD49a^-^Eomes^-^ cNKs accumulated in metastatic livers in *Hobit*^-/-^ but not in *Nkp46*^iCre^*Eomes*^fl/fl^ mice, which confirms their cNK cell origin (Figures 5E-G). Also, the blood of mice with LLC liver metastasis contained increased numbers of CD49a^-^Eomes^-^ cNKs, suggesting that LLC metastases induce systemic changes in the cNK compartment (Supplementary Figure S4D). In MC38-derived metastatic nodules, we detected another unique subset of cNKs. Despite the expression of immature markers CD69, Thy1.2 and CD27, this subset expressed characteristic markers of differentiation (T-bet, Eomes and GrzB) as well as the integrin CD49a (Figures 5H-K). We refer to these cells as CD49a^+^Eomes^+^ NK cells, and show that also this subset arose from cNKs, since it was only found in metastatic livers of *Hobit*^-/-^ but not in those of *Nkp46*^iCre^*Eomes*^fl/fl^ mice (Figures 5L-N). In contrast to mice with LLC-derived liver metastasis, we did not observe high frequencies of CD49a^+^Eomes^+^ cNKs in the blood of mice with MC38-derived metastases, although the systemic population characteristic of this tumor type resembles the less differentiated cNK_2 cluster (Supplementary Figure S4E). The emergence of CD49a^-^Eomes^-^ cNKs was not a unique feature of the LLC tumor type or C57BL/6 mice, since we found similar CD49a^-^Eomes^-^ cNKs in livers of BALB/c mice with 4T1-derived metastases (Supplementary Figures S5A-S5F). Along the same lines, CD49a^+^Eomes^+^ cNKs were not unique to MC38 liver metastases or C57BL/6 mice, as we also found them in BALB/c mice with CT26-derived liver metastases (Supplementary Figures S5G-S5L). These findings suggest a dichotomy of cancers that impair cNKs (LLC, 4T1) and those that do not (MC38, CT26). It was recently described that cancer-cell-intrinsic cGAS supports NK-mediated control of cancer (Marcus et al., 2018). Because MC38 and CT26 express cGAS in contrast with LLC and 4T1 (Schadt et al., 2019), we analyzed whether cancer-cell-intrinsic cGAS may be one of the factors driving the emergence of specific cNK subsets. Therefore, we overexpressed cGAS in LLC and found that this resulted in a reduced metastatic load in the liver (Supplementary Figure S5M) and a reduction of the proportion of Eomes^-^CD49a^-^ cNKs (Supplementary Figure S5N).

**Figure 5:**
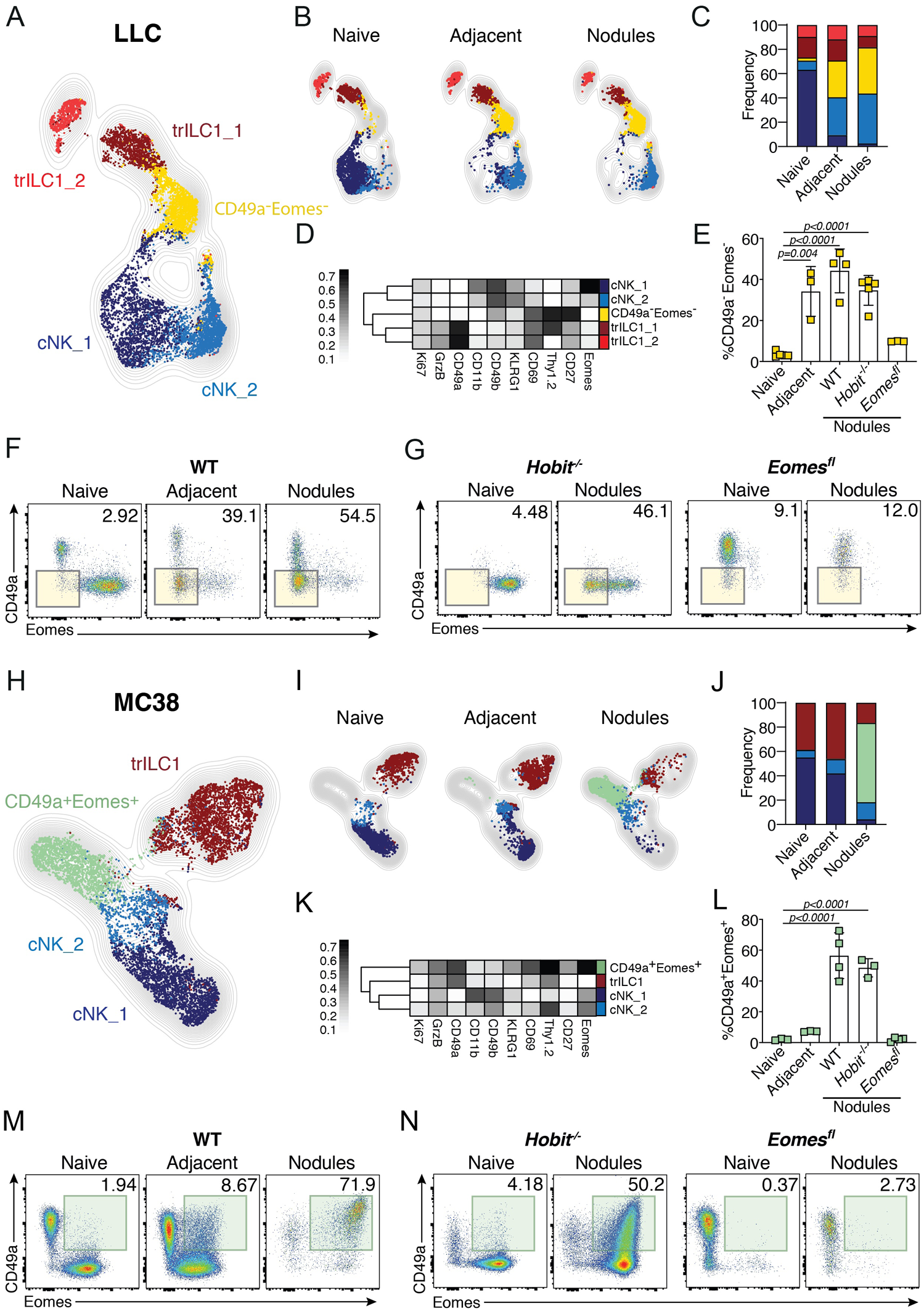
Metastatic livers drive the differentiation of unique cNK populations. NK cells from naïve and day 21 metastatic (nodules and adjacent tissue) livers were analyzed by multi-parameter single-cell mapping using flow cytometry. **(A-G)** LLC-metastatic and control livers. **(H-N)** MC38-metastatic and control livers. Samples were pre-gated on single live CD45^+^lineage^-^ cells and subsequently gated on NK1.1^+^NKp46^+^ cells. cNK = conventional NK cells, CD49a^-^CD49b^+^ ; trILC1 = tissue-resident ILC1s, CD49a^+^CD49b^-^. **(A, H)** UMAP projections overlaid with FlowSOM-guided manual metaclusters displaying cNKs and trILC1s from all samples. DP = CD49a^+^Eomes^+^. **(B, I)** UMAP projections overlaid with FlowSOM-guided manual metaclusters separated by sample category (naïve, adjacent, nodules). **(C, J)** Relative frequency of each cluster in the different sample categories (naïve, adjacent, nodules). **(D, K)** Heatmap summary of median marker expression values of the different markers analyzed for each cluster. **(E, L)** The frequency of unique metastasis-induced subsets. **(E)** CD49a^-^Eomes^-^ NK cells in LLC-metastasis. **(L)** CD49a^+^Eomes^+^ NK cells in MC38-metastasis. The bar represents the mean ± SD, symbols represent livers from individual mice. One-way analysis of variance (ANOVA), with Tukey’s multiple comparisons test. Experiments were performed at least twice with similar results. **(F, M)** Representative dot plots of cNKs and trLC1s cells for each sample category based on their expression of CD49a and Eomes. **(F)** LLC-metastasis. Highlighted in yellow is the CD49a^-^Eomes^-^ population observed in LLC adjacent tissue and nodules. **(M)** MC38-metastasis. Highlighted in green is the CD49a^+^Eomes^+^ population observed in MC38 nodules. **(G, N)** Representative dot plots of cNKs and trILC1s isolated from naïve liver or metastatic nodules from *Hobit*^-/-^ and *Eomes*^*fl*^ mice, based on their expression of CD49a and Eomes. **(G)** LLC-metastasis. Highlighted in yellow is the CD49a^-^Eomes^-^ population observed in LLC adjacent tissue and nodules. **(N)** MC38-metastasis. Highlighted in green is the CD49a^+^Eomes^+^ population observed in MC38 nodules. Groups contained 3-5 mice. Experiments were performed at least twice with similar results.

Our high dimensional analyses also confirmed that trILC1s were mostly located outside the metastatic nodules (Figures 5B, 5C, 5I and 5J), as previously revealed by immunofluorescence. In line with their preserved cytotoxicity, trILC1s only underwent slight phenotypic changes in the nodules and their adjacent tissue of metastatic livers from both LLC and MC38 origin (Figures 5B, 5C, 5I and 5J).

Altogether, we uncovered unique populations of cNKs in metastatic livers, and found that cancer-specific features drive the differentiation of these cells. Thus, whereas certain tumor types (e.g. LLC and 4T1) promote the emergence of very immature subsets of cNKs, others such as MC38 or CT26 bear a unique intranodular cNK cell subset that retains key features of differentiation.

### Single-cell RNA-sequencing reveals novel transcriptional signatures of cNKs populating the hepatic metastatic niche

To gain further insights into the nature of these unique and previously undiscovered cNK cell subsets populating the hepatic metastatic niche, we performed 10x single-cell RNA-seq on CD45^+^ lin^-^NK1.1^+^NKp46^+^ NK cells sorted from naïve as well as from MC38 and LLC metastatic livers. The clusters were subsequently projected on a UMAP plot with each dot representing a single cell. The identity of cNK and trILC1 cells was confirmed by their uniformly high expression of the lineage marker gene *Ncr1* (Nkp46), as well as the expression of *Eomes* (Eomes, for cNKs) and *Itga1* (CD49a, for trILC1s) (Supplementary Figure S6A).

We identified two clusters for trILC1s and 6 clusters corresponding to cNKs (Figure 6A). The top genes identifying each cluster are displayed in Supplementary Table 1 and in a heatmap (Supplementary Figure S6B). We found that cNK_4 and cNK_6 were preferentially enriched in LLC-derived and MC38-derived metastases, respectively (Figures 6B and 6C), while the cNK_5 cluster expanded in either environment (Figures 6B and 6C).

**Figure 6:**
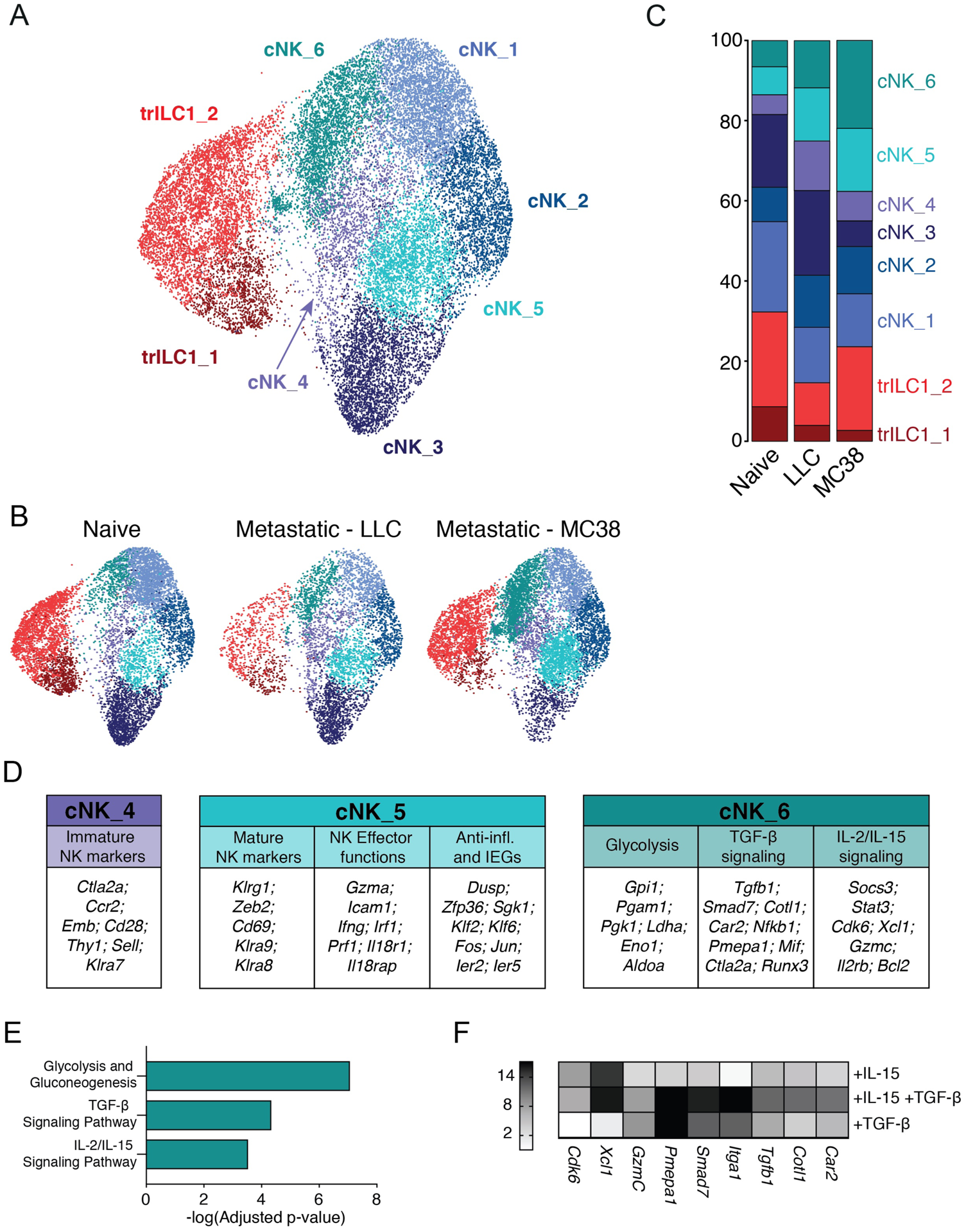
Single-cell RNA-sequencing reveals novel transcriptional signatures of cNKs in the hepatic metastatic niche. Single-cell RNA-sequencing was performed on NK cells sorted from naïve, LLC-metastatic and MC38-metastatic livers (6 mice per condition, pooled into 1 sample for droplet encapsulation and library preparation). **(A-C)** UMAP projections identifying the different cNK/trILC1 clusters in **(A)** all samples or **(B)** individual samples with **(C)** bar plots showing the relative frequency of the different clusters in each sample. **(D)** Top upregulated genes in cluster cNK_4, cNK_5 and cNK_6 arranged in functional categories. **(E)** Pathway analysis of significantly upregulated genes in cluster cNK_6. **(F)** cNKs and trILC1s were sorted from naïve livers and cultured *in vitro* for 48 h with 25 ng/ml mouse IL-15/IL-15R complex and/or 10 ng/ml human TGFβ1. Transcripts of cNK_6 cluster-specific genes were quantified by qPCR. The heatmap shows the average fold change relative to the untreated condition. The data represent 3 biological and 3 technical replicates, and the experiment was performed 3 times with similar results.

The majority of differentially expressed genes (DEGs) from the cNK_4 cluster suggested an immature phenotype, as illustrated by the high expression of *Ctla2a, Ccr2, Emb, Cd28, Thy1, Sell* (CD62L) or *Klra7* (Ly49G2) and the strong downregulation of *Gzmb* (Supplementary Table 1, Figures 6C and 6D). Also on the protein level, we found an upregulation of Thy1.2, CD62L and Ly49G2 in cNKs from MC38- and LLC-derived metastatic nodules (Supplementary Figures S6C and S6D). Further, the transcriptional signature of this cluster showed a striking resemblance with that of an immature NK population previously described in steady state spleen (Crinier et al., 2018). In contrast, NK cells from the cNK_5 cluster, common to both LLC and MC38 metastatic livers, appeared to be more differentiated expressing *Klrg1, Cd69*, as well as *Klra9* (Ly49I) and *Klra8* (Ly49H) (Figure 6D and Supplementary Table 1). This cluster is characterized by high expression of effector genes (*Icam1, Ifng, Gzma* and *Prf1)*, a few genes involved in anti-inflammatory responses (*Dusp1, Zfp36* or *Sgk1*) and several Immediate Early Genes (IEGs) (*Fos, Jun, Klf2, Klf6, Ier2* and *Ier5*) (Figure 6D and Supplementary Table 1). The latter may indicate an “alertness” of cells in cluster cNK_5, allowing them to rapidly respond to environmental cues. Additionally, the cNK_5 cluster strongly resembled the cNK_3 cluster enriched in naïve livers and underrepresented in metastases (Supplementary Figure S6E). Cells from the cNK_5 cluster however expressed higher amounts of cytotoxic genes (*Prf1* or *Gzma)* and genes from the IL-18 pathway (*Il18r1, Il18rap*), suggesting its skewing towards a more effector phenotype.

cNK_6 was the only cNK cell cluster expressing *Itga1*, increased amounts of *Thy1* (Supplementary Figure S6F) and *Eomes* (data not shown) transcripts compared to other clusters and thus showed the highest similarity to the CD49a^+^Eomes^+^ cNK subset identified by flow cytometry. About half of the genes that define the immature cNK_4 cluster were highly expressed by cells of the cNK_6 cluster (Figure 6D, Supplementary Table 1 and Supplementary Figure S6G). We found an increased expression of several TGF-β-induced genes such as *Smad7, Runx3, Pmepa1, Plac8, Car2, Cotl1* and *Tgfb1* itself (Figure 6D and Supplementary Table 1). In agreement, EnrichR pathway analysis identified upregulation of the “TGF-β Signaling Pathway” in the cNK_6 cluster (Figure 6E) (Chen et al., 2013). This analysis also revealed an enrichment of a gene set related to the “IL-2/IL-15 Signaling Pathway” (*Gzmc, Socs3, Stat3, Il2rb, Bcl2, Cdk6, Xcl1*) (Figures 6D and 6E) and another gene set related to “Glycolysis and Gluconeogenesis” (*Ldha, Eno1, Pgam1, Gpi1, Pgk1*) (Figures 6D and 6E). The latter pathway can be induced by the synergy of hypoxia and IL-15 (Squez et al., 2016). We validated the importance of the TGF-β and IL-15 signatures in cNK_6 by assessing the expression of selected DEGs upon *ex vivo* stimulation of isolated cNKs with TGF-β and/or IL-15 (Figure 6F).

Taken together, whereas the transcriptome of cNK_4 cells expanded in LLC-derived metastases is reminiscent of an immature status, that of cNK_6 cells from MC38-derived metastases appears to be the result of dual TGF-β and IL-15 signaling. This illustrates that tumors can employ differential mechanisms to drive NK cell phenotype and function.

### IL-15 and TGF-β modulate the fate of CD49a^+^Eomes^+^ cNKs in metastatic nodules

Our discovery that cNKs populating liver metastases have a unique signature suggests that the metastatic microenvironment regulates the differentiation of cNKs. Furthermore, it appears that each tumor influences cNKs differently, presumably dependent on specific features associated with individual tumors. Because the single-cell RNA-sequencing data pointed towards the TGF-β and IL-15 pathways, we quantified the transcripts for these cytokines in the hepatic metastatic niche. We found high amounts of *Tgfb1* in the adjacent tissue and nodules of both MC38- and LLC-metastatic livers (Figure 7A). We detected low amounts of transcripts of NK cell-modulating cytokines such as *Il2, Il12* and *Il18* in metastatic livers from both tumor types (data not shown). In contrast, we found high amounts of transcripts of *Il15* and *Il15ra* in MC38-but not LLC-nodules (Figures 7B and 7C), which may explain that MC38-nodules contained significantly more cNKs than MC38-nodule-adjacent tissue, LLC nodules or naïve liver (Figures 1E-G).

**Figure 7:**
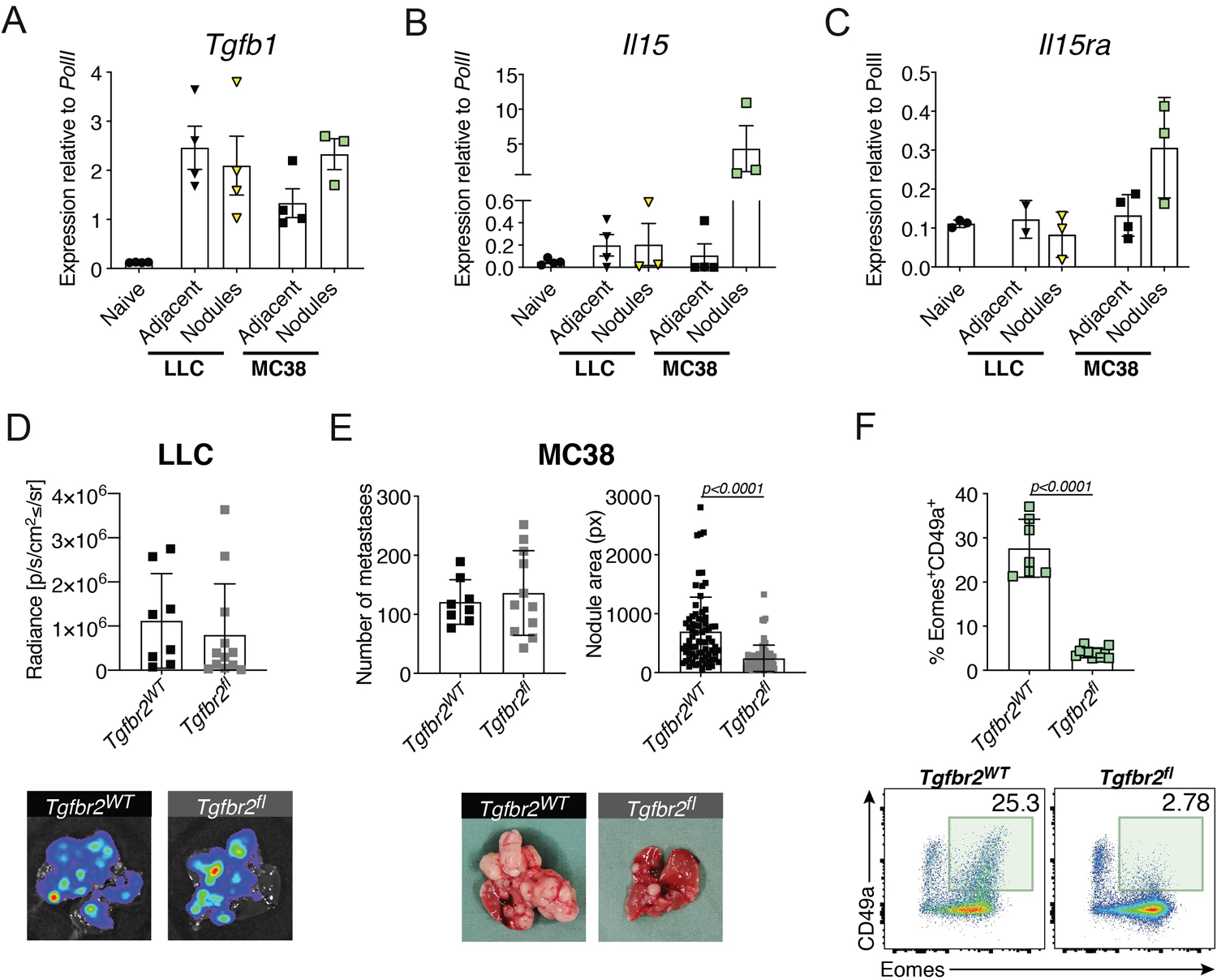
IL-15 and TGF-β modulate the fate of CD49a^+^Eomes^+^ cNKs in metastatic nodules. **(A, B)** Quantification of transcripts of **(A)** *Tgfb1*, **(B)** *II15*, and **(C)** *II15ra* in lysates of naïve, LLC- and MC38-metastatic livers. Bars show the mean ± SD. Each symbol represents an individual mouse. One-way analysis of variance (ANOVA), with Tukey’s multiple comparisons test. The experiment was performed twice with similar results. **(D)** Upper panel: Quantification of LLC liver nodules from *Tgfbr2*^WT^ and *Tgfbr2*^fl^ mice by *ex vivo* IVIS 21 days after tumor cell injection. Bars show the mean ± SD. Each symbol represents an individual mouse. One-way analysis of variance (ANOVA), with Tukey’s multiple comparisons test. The experiment was performed twice with similar results. Lower panels: Representative IVIS images of metastatic livers from each group at the endpoint. *Tgfbr2*^*WT*^ *= Ncr1*^*iCre/wt*^.*Tgfbr2*^*wt/wt*^ ; *Tgfbr2*^*fl*^ *= Ncr1*^*iCre/wt*^.*Tgfbr2*^*fl/fl*^. **(E)** Upper panel: Quantification of macroscopic MC38 liver nodules (number and area) from *Tgfbr2*^*WT*^ and *Tgfbr2*^*fl*^ mice 21 days after tumor cell injection. Bars show the mean ± SD. Each symbol represents an individual mouse. One-way analysis of variance (ANOVA), with Tukey’s multiple comparisons test. The experiment was performed twice with similar results. Lower panels: Representative images of metastatic livers from each group at the endpoint. *Tgfbr2*^*WT*^ *= Ncr1*^*iCre/wt*^.*Tgfbr2*^*wt/wt*^ ; *Tgfbr2*^*fl*^ *= Ncr1*^*iCre/wt*^.*Tgfbr2*^*fl/fl*^. **(F)** Upper panel: Percentage of CD49a^+^Eomes^+^ NK cells in MC38 nodules from *Tgfbr2*^*WT*^ and *Tgfbr2*^*fl*^ mice. Bars show the mean ± SD. Each symbol represents an individual mouse. One-way analysis of variance (ANOVA), with Tukey’s multiple comparisons test. The experiment was performed twice with similar results. Lower panel: Representative dot plots of cNKs and trILC1s in MC38 nodules from *Tgfbr2*^*WT*^ and *Tgfbr2*^*fl*^ mice, with the CD49a^+^Eomes^+^ population highlighted in green.

We then crossed *Ncr1*^iCre^ to *Tgfbr2*^fl/fl^ mice to specifically delete TGF-β signaling in NK cells. Despite the previously reported TGF-β-mediated inhibition of NK maturation during their development (Marcoe et al., 2012), we did not observe phenotypical differences in cNKs or trILC1s of *Ncr1*^iCre^*Tgfbr2*^fl/fl^ mice compared to control mice in steady state (Supplementary Figures S7A-S7C). The LLC-derived metastatic load as well as the phenotype of cNKs and trILC1s in *Ncr1*^iCre^*Tgfbr2*^fl/fl^ mice were similar to those in control mice (Figure 7D and Supplementary Figure S7D). However, hepatic MC38-derived metastases were significantly reduced in mice lacking the TGF-β receptor on NK cells (Figure 7E). Although the NK cell population did not undergo major phenotypic changes (Supplementary Figure S7E), the CD49a^+^Eomes^+^ cNK cell subset infiltrating the metastatic nodules in control mice failed to develop in *Ncr1*^iCre^*Tgfbr2*^fl/fl^ mice (Figure 7F).

Taken together, cytokines produced in the metastatic niche influence the phenotype and function of tumor-associated NK cells in an individual fashion as exemplified here by IL-15 and TGF-β in case of LLC and MC38 liver metastasis. Unraveling such pathways may lead to discovery of druggable targets and improved NK cell-based therapies for metastatic disease.

## DISCUSSION

The liver is a preferred metastatic site for many different cancer types (Williamson et al., 2019), suggesting an exceptional immune environment. Indeed, the liver is equipped with mechanisms that promote immune tolerance of CD8^+^ T cells (Limmer et al., 2000; Zheng and Tian, 2019) to prevent deleterious immune responses against innocuous antigens. In addition, the liver is surveilled by passer-by cNKs and by resident trILC1s with distinct features (Peng and Tian, 2017). These abundant innate lymphocytes may be the major effectors in hepatic immune defense.

In the context of viral infection, trILC1s suppressed T cell-mediated viral control (Zhou et al., 2019) and were protective against acute liver injury (Nabekura et al., 2020), suggesting a regulatory rather than effector function. Using genetically modified mice that lack cNKs or trILC1s, we discovered that both innate subsets collaborated in a non-redundant fashion to control hepatic metastasis. Specifically, we found that trILC1s mainly interfered with seeding of cancer cells, whilst cNKs controlled metastatic outgrowth and progression. The doorkeeper function of trILC1s may be explained by their unique location in the hepatic sinusoids (Peng et al., 2013). A significant proportion of trILC1s express CXCR6, which may mediate retention to CXCL16-producing sinusoidal cells (Paust et al., 2010). Our observation that trILC1s hardly infiltrated metastatic nodules is in line with their inability to control cancer cells beyond seeding. At the same time, keeping their distance from disseminated cancer cells may explain why trILC1s sustain their cytotoxic function even at a late stage of metastatic disease.

The limited capacity to infiltrate metastatic nodules is also a hallmark of cNKs arising in tumors of LLC origin. The cNK cell subset instructed by this tumor type expresses low amounts of CD49b, Eomes and T-bet, suggesting stalled maturation (Gordon et al., 2012) or exhaustion (Gill et al., 2012). Along the same lines, we found that cNKs from LLC-metastatic livers were unresponsive to a variety of stimuli. In contrast, CD49a^+^CD49b^+^Eomes^+^ cNKs readily infiltrated MC38-derived metastatic nodules and maintained several of their functions. The expression of CD49a and additional immature markers (Thy1.2, CD69, CD27) associated with this population is reminiscent of an intratumoral ILC1-like subset (CD49a^+^CD49b^−^Eomes^int/low^) reported in methylcholanthrene-induced sarcomas (Gao et al., 2017). However, the cNK subset we found in MC38-metastatic nodules may rather represent transiently activated immature cNK cells, because of their high expression of differentiation markers such as T-bet, CD49b and Eomes. A CD49a^+^CD49b^+^ NK cell population was described in the liver following viral infection, although with a more mature phenotype (Li et al., 2020).

The differences between cNKs associated with LLC- or MC38-derived metastasis reveal that disseminated cancer cells orchestrate the metastatic niche in a cancer-specific fashion, influencing the phenotype and function of cNKs. For example, we detected a high expression of the survival factor IL-15 (Ranson et al., 2003) in MC38-but not LLC-derived metastasis, which may explain the paucity of cNKs in the latter tumor type. Further, we detected considerable amounts of TGF-β in both MC38- and LLC-metastatic livers. Because TGF-β impairs NK cell cytotoxicity and maturation (Lee et al., 2004; Marcoe et al., 2012; Viel et al., 2016), we investigated metastatic control in *Nkp46*^iCre^*Tgfbr2*^fl/fl^ mice in which NK cells can’t respond to TGF-β. In such mice, cNK cells did not convert into CD49a^+^Eomes^+^ cNKs and more importantly, the MC38-derived metastatic load was significantly reduced. Thus, TGF-β directly regulates the expression of CD49a, which is agreement with previous reports (Gao et al., 2017; Rautela et al., 2019), and moreover, directly compromises the anti-metastatic efficacy of cNK cells. Despite the presence of TGF-β in LLC-metastases, preventing TGF-β-signaling did not change the LLC-metastatic load, suggesting that additional mechanisms limit anti-metastatic cNK functions in LLC metastasis. Alternatively, TGF-β blockade may only be efficient when cNK cells face a metastatic microenvironment that favors their survival, e.g. in the presence of IL-15.

We found that the cNK cell subsets present in the metastatic niche of LLC or MC38 also emerged in hepatic metastases derived from other cell lines. Indeed, CT26 influenced cNKs in a similar way as MC38 did, whereas the impact of 4T1 resembled that of LLC. These findings point towards a dichotomy, which is reminiscent of the concept in which cancers are described as hot or cold, a classification mainly based on the CD8^+^ T cell landscape within a tumor (Galon et al., 2006). Recently, a four-category classification was proposed – hot, altered-excluded, altered-suppressed and cold – which considers other parameters as well (Galon and Bruni, 2019). The relevance of such classifications for the quality of innate immune defense is not well studied but it is likely that at least some factors promoting T cell immunity also support innate lymphoid cells. As discussed above, one such factor may be IL-15 (Burkett et al., 2004; Ranson et al., 2003; Sato et al., 2007). IL-15 is induced by the cancer-cell-intrinsic factor cGAS through the activation of the stimulator of interferon genes (STING) pathway (Nicolai et al., 2020; Santana Carrero et al., 2019). STING activation in myeloid cells induces the production of type I IFN, which supports NK and CD8^+^ T cells responses directly or indirectly through the production of IL-15 (Marcus et al., 2018; Nicolai et al., 2020; Schadt et al., 2019). Interestingly, hot cancers display a type I IFN signature (Gajewski et al., 2013) and both MC38 and CT26 cells contain high amounts of active cGAS and produce spontaneous type I IFN themselves (MC38) or induce its production in adjacent myeloid cells (CT26), whereas LLC and 4T1 do not (Schadt et al., 2019).

Taken together, our study demonstrates collaborative roles of circulating and tissue resident innate lymphocytes combatting the formation and growth of metastasis. The unique properties of the cNK subsets described here and their differentiation are instructed by the metastatic microenvironment. Thus, understanding the regulatory hierarchy of metastatic-derived factors for each individual tumor type will be crucial to exploit the anti-metastatic potential of innate lymphoid cells in metastasis.

## ACKNOWLEDGEMENTS

We thank Karina Silina, Mirjam Lutz, Virginia Cecconi (Institute of Experimental Immunology, University of Zurich), the Flow Cytometry Facility (University of Zurich) and Emilio Yangüez and Ge Tan from the Functional Genomics Center Zurich for assistance. This work was supported by the Swiss National Science Foundation (CRSII5_177208 and 310030_175565 to MvdB, PR00P3_179775 to ST, 316030_150768 and 310030_146130 to BB), the Swiss Cancer league (KLS-4098-02-2017 to MvdB, KFS-4431-02-2018 to ST and BB), the Promedica Stiftung (ST), the University of Zurich (University Research Priority Project “Translational Cancer Research” to MvdB and BB), and the Monique-Dornonville-de-la-Cour Foundation Zurich (MvdB).

## AUTHOR CONTRIBUTIONS

G.Lu., L.D., M.vdB. and S.T. designed the research. A.M., D.B., G.Li., G.Lu., J.G., L.D., L.H., L.W., M.N., M.V. and N.N. performed research and analyzed data. B.B., G.Lu., L.D., M.vdB. and S.T. wrote the manuscript.

## DECLARATION OF INTERESTS

The authors declare no competing interests.

## STAR METHODS

### Mice

Female 6-10-weeks-old C57BL/6 mice were obtained from Janvier Labs (Roubaix, FR). *Ncr1*^Cre/wt^ mice (B6.Ncr1^tm1.1(icre)Viv^) were provided by Eric Vivier (Marseille, France), *Rosa26*^iDTR^ mice by Ari Waisman (Mainz, Germany); *Eomes*^fl/fl^, *Ai14*^fl/fl^ and *Cd1d*^-/-^ mice were obtained from the Jackson Laboratory. *Nkp46*^iCre^ mice were crossed to *Rosa26*^iDTR^ mice to obtain *Nkp46*^iCre/wt^ *Rosa26*^iDTR/wt^ (*Nkp46*^iCre^*R26R*^*i*DTR^) or *NKp46*^iCre/wt^*Rosa26*^wt/wt^ controls (*Nkp46*^iCre)^. Mice homozygous for the *Hobit* targeted mutation (*Hobit*^-/-^ mice) were provided by Klaas van Gisbergen (Mackay et al., 2016). *B6*.*129-Nfil3(loxP)tm1Rbrc* (*Nfil3*^*fl/fl*^) mice were generated by Motomura Y. (Motomura et al., 2011) and provided by Henrique Veiga-Fernandes, Champalimaud Foundation, Lisbon, Portugal. Mice were housed in the Laboratory Animal Services Center (LASC) of the University of Zurich according to institutional guidelines under specific pathogen-free conditions. All animal experiments were approved by the Swiss cantonal veterinary office (license numbers 65/2015 and 156/2018).

### Cell lines

MC38 cells and 4T1 cells were provided by Carlotta Tacconi and Michael Detmar (Institute of Pharmaceutical Sciences, ETH, Zurich, Switzerland), respectively. Lewis Lung carcinoma (LLC) and CT26 colon carcinoma cells were purchased from ATCC. LLC cells were lentivirally transduced to express firefly luciferase of cGAS to generate LLC-LUC or LLC-cGAS (Schadt et al., 2019) cells, respectively. Viral particles were a gift from Christian Münz (University of Zurich). All cancer cells were cultured in Dulbecco’s modified Eagle’s medium (DMEM, GIBCO) supplemented with 10% fetal calf serum (FCS, ThermoFisher Scientific), 30 U/ml Penicillin, 30 µg/ml Streptomycin (antibiotics, ThermoFisher Scientific) and 2mM L-Glutamine (ThermoFisher Scientific). All cancer cell lines were used between passages 2 and 10. Cell lines were confirmed to be free of *Mycoplasma* ssp. by PCR and various viruses by Charles River Research Animal Diagnostic Services.

### Model of experimental liver metastasis

To generate murine liver metastases, 6-10-weeks-old mice were anesthetized using vaporized isoflurane, and the upper lateral abdominal wall was incised followed by the injection of 5×10^5^ MC38, or 2×10^5^ LLC-LUC, CT26 or 4T1 cells in 40 µl PBS into the splenic parenchyma. Immediately after injection, the spleen was excised to prevent growth of splenic tumors. The wound was closed using sutures (70 cm Coated Vicryl, ETHICON) and autoclips (Autoclip 9 mm, Becton Dickinson). Control mice underwent mock surgery. Livers were analyzed 21-28 d after tumor cell injection. MC38-derived metastatic nodules were counted using a stereotactic microscope and the nodule area was quantified using the ImageJ software, whereas LLC-derived metastasis were visualized and quantified using an IVIS 200 imaging system (PerkinElmer) 20 min after i.p. injection of 150 mg/kg D-Luciferin (Promega). The data are presented as total luminescence per liver for each mouse.

### Depletion of cells by antibodies or diphtheria toxin (DTX)

Mice were treated i.p. with 50 µl of anti-asialo GM1 antibodies (Wako Pure Chemical Industries) or 0.2 mg rabbit IgG (Sigma) twice a week starting 48 h before intrasplenic injection of tumor cells. Diphtheria toxin from *Corynebacterium diphtheria* (Calbiochem, 250 ng/mouse for initial depletion and 125 ng/mouse for the following injections) diluted in PBS was injected i.p. at days -2, -1, 2, 5, 8, 11, 15 and 18 relatively to tumor injection. Anti-CD8α (100 µg/mouse, clone YTS169.4, BioXCell) was injected i.p. 2 days before tumor injection. Depletion efficiency was monitored in the blood by flow cytometry.

### Flow cytometry

Livers were harvested in RPMI supplemented with 10% FCS, digested with Collagenase IV (1 mg/ml, Bioconcept) and DNaseI (2.6 μg/ml, ThermoFisher Scientific) for 45 minutes at 37°C. Cells were washed with PBS and filtered through a 100-μm filter to remove debris. Lymphocytes were purified using a density gradient centrifugation step (Percoll, GE Healthcare). Blood was collected in PBS containing 2 mM EDTA. Red blood cells were lysed in all samples using RBC lysis buffer (4.15 g NH4Cl, 0.55 g KHCO3, 0.185 g EDTA in 500 ml ddH2O) for 1.5 minutes.

Single cells were incubated for 15 minutes in 250 µg/ml anti-mouse CD16/32 (clone 2.4G2) in PBS to block Fc-receptors, washed with PBS, and surfaced-stained in 50 μl antibody-mix in PBS. For intracellular cytokine staining, cells were stimulated with 100 ng/ml phorbol 12-myristate 13-acetate (PMA) plus 1 µg/ml ionomycin, 100 ng/ml recombinant mouse IL-18 (MBL International) plus 10 ng/ml recombinant mouse IL-12 p70 (Peprotech), 10 ng/ml recombinant mouse IL-15 (Peprotech) or anti-NK1.1 (clone PK136, BioXCell) for 4 h at 37°C in the presence of GolgiPlug™/GolgiStop™ (BD Pharmigen™). Cells were stained for surface molecules as described above, washed with PBS, and fixed for 60 min on ice using IC Fixation Buffer from Foxp3/Transcription Factor Staining Buffer Set (eBioscience). Subsequently, cells were stained for intracellular cytokines in 50 μl permeabilization buffer from the Foxp3/Transcription Factor Staining Buffer Set overnight at 4°C. After washing with permeabilization buffer, samples were suspended in FACS buffer (PBS, 20 mM EDTA pH 8.0, 2% FCS) and acquired FACS LSRII Fortessa (BD Bioscience) or FACSymphony (BD Biosciences).

To determine cell numbers, CountBright™ Absolute Counting Beads were used (ThermoFisher Scientific). Dead cells were excluded using Live/Dead fixable staining reagents (Invitrogen), and doublets were excluded by FSC-A/FSC-H gating. Antibodies: CD115-Biotin (AFS98; BioLegend), CD19-Biotin (6D5; BioLegend), CD3e-Biotin (145-2C11; BioLegend), CD5-Biotin (53-7.3; BioLegend), Ly6G-Biotin (1A8; BioLegend), NK1.1-FITC (PK136; BioLegend), NK1.1-BV711 (PK136; BioLegend), NK1.1-BV785 (PK136; BioLegend), NKp46-PerCP-eFluor710 (29A1.4; eBioscience), NKp46-FITC (29A1.4; eBioscience), CD4-BV785 (104 (SJL); BioLegend), CD45-APC-Cy7 (30-F11, BioLegend), CD49a-BV510 (Ha31/8; BD Biosciences), CD49a-BUV395 (Ha31/8; BD Biosciences), CD49b-PB (DX5; BioLegend), CD27-PE (LG.3A10; BioLegend), CD27-PE-Cy7 (LG.3A10; BioLegend), CD62L-BUV737 (MEL-14; BD Biosciences), CD69-BV605 (H1.2F3; BioLegend), Thy1.2-BV785 (30-H12; BioLegend), Thy1.2-AF700 (30-H12; BioLegend), KLRG1-BV510 (2F1/KLRG1; BioLegend), CD11b-BV711 (M1/70; BioLegend), CD11b-BUV737 (M1/70; BioLegend), Ly49G2-PerCP-eFluor710 (4D11; eBioscience), EOMES-PE-texRed (Dan11mag; eBioscience), GrzB-AF647 (GB11; BioLegend), Ki67-PerCP-eFluor710 (SolA15; eBioscience), Tbet-PE-Cy7 (4B10; BioLegend), IFN-γ-PE-Cy7 (XMG1.2; BioLegend), CD107a-APC (1D4B; BioLegend), CD8a-BV605 (53-6.7; Biolegend), Streptavidin-APC (BD Biosciences), Streptavidin-BUV563 (BD Biosciences), Zombie NIR™ Fixable Viability Kit (BioLegend), Zombie UV™ Fixable Viability Kit (BioLegend).

### Analysis of flow cytometry data

Flow cytometry data were preprocessed, down-sampled and exported using FlowJo v10 software (Tree Star). The exported FCS files were uploaded in Rstudio (R software environment, version 3.4.0) and were transformed with an inverse hyperbolic sine (Arsinh) function. UMAP projections were calculated and FlowSOM-based clustering was performed using the guided manual metaclustering (Brummelman et al., 2019; Nowicka et al., 2017). All plots were drawn with ggplot2.

### Single-cell RNA sequencing using 10x Genomics platform

NK cells (live CD45+CD3-CD5-Ly6G-CD19-CD115-NK1.1+Nkp46+) were sorted from naïve, LLC and MC38 metastatic livers. For each sample, cells were pooled from 6 biological replicates. The quality and concentration of the single cell preparations were evaluated using an hemocytometer in a Leica MD IL LED microscope and adjusted to 1’000 cells/ml. 10’000 cells per sample were loaded in to the 10X Chromium controller and library preparation was performed according to the manufacturer’s indications (Chromium Next GEM Single Cell 3’ Reagent Kits v3 protocol). The resulting libraries were sequenced in an Illumina NovaSeq6000 sequencer according to 10X Genomics recommendations (paired-end reads, R1=28, i7=8, R2=91) to a depth of around 50,000 reads per cell. FASTQ files were created with the Cell Ranger demux pipeline. Reads were pseudo-aligned to a mouse reference (ensemble release 97) with the kallisto/bustools pipeline, using kallisto v0.46.0 (Bray et al., 2016) and bustools <u>v0.39.3</u> (Melsted et al., 2019). The resulting count matrix was analyzed with Seurat v3 (Stuart et al., 2019). Data were scaled and transformed using SCTransform v0.2.0 (Hafemeister and Satija, 2019) for variance stabilization. The experiment resulted in the following number of obtained cells: 9164 cells from naïve livers (median of 1543 unique genes detected per cell), 8910 cells from LLC metastatic livers (median of 1681 unique genes detected per cell), and 9565 cells from MC38 metastatic livers (median of 1606 unique genes detected per cell). Contaminating cell clusters that were NKp46 negative and did not consist of NK cells were excluded from further analyses. Principle Component Analysis was performed on the expression of the detected variable genes. The first 50 principal components were included for further downstream analyses based on visual inspection of Seurat’s PCElbowPlot. All cells were clustered based on the principal component analysis with the Louvain algorithm using the following granularity parameters: resolution = 0.5. Differential marker expression analyses were conducted with the Seurat FindMarkers and FindAllMarkers functions. 10x libraries were prepared and sequenced at the Functional Genomics Center (University of Zürich).

### Gene ontology analysis

Gene Ontology (GO) analysis was performed with EnrichR (Chen et al., 2013). Venn diagrams were created using the jvenn online tool (Bardou et al., 2014).

### Degranulation assay and IFN-γ production

Cell suspensions were cultured in Roswell Park Memorial Institute (RPMI, GIBCO) supplemented with 10% FCS (ThermoFisher Scientific) and 2mM L-Glutamine (ThermoFisher Scientific) with either 100 ng/ml PMA plus 1 µg/ml ionomycin, 100 ng/ml IL-18 (MBL) plus 10 ng/ml IL-12 p70 (Peprotech), 10 ng/ml IL-15 (Peprotech) or plate-bound anti-NK1.1 (clone PK136, BioXCell) for 1 h in 96-well plates at 37°C. CD107a antibody (1:200 final dilution, 1D4B BioLegend) was added to the mix. Subsequently, GolgiPlug™ (10 µg/ml, BD Pharmigen™) and GolgiStop™ (10 µg/ml, BD Pharmigen™) were added and cells were incubated for additional 3 h. For anti-NK1.1 activation 96 well plates were coated ON at 4°C with 10 µg/ml anti-NK1.1 antibody (PK136) or isotype control (C1.18.4), washed with PBS and blocked with 20% FCS for 20 min. After incubation cells were collected and analyzed by flow cytometry.

### Killing assay

Liver NK cells (live, CD45^+^CD3^-^CD5^-^Ly6G^-^CD19^-^CD115^-^ NK1.1^+^NKp46^+^CD49a^+^CD49b^-^ and CD45^+^CD3^-^CD5^-^Ly6G^-^CD19^-^CD115^-^ NK1.1^+^NKp46^+^CD49a^-^CD49b^+^) were sorted with a BD FACSAria™ III sorter. Target cells (MC38, LLC) were stained with PKH26 Red Fluorescent Cell Linker Mini Kit (Sigma) following manufacturer’s instructions, and seeded in 96-well round-bottom plates with 10^4^ cells/well. NK cells were incubated with target cells at effector:target ratios of 1:1, 4:1 and 8:1 for 24 h at 37°C in 5% CO2. After removal of medium, 0.8 µM TO-PRO-3 (ThermoFisher Scientific) was added, and cells were acquired on a LSRII Fortessa flow cytometer (BD). Data were analyzed using FlowJo v10 software (Tree Star). Percentage specific killing was calculated as follows: [(% cell death in the presence of NK cells) — (% spontaneous cell death)] ≑ (100 — % spontaneous cell death).

### Quantitative real-time PCR

Metastatic livers were divided in adjacent tissue and nodules. RNA was isolated from lysates of naïve or metastatic livers from C57BL/6 mice using the Qiagen Micro Kit according to the manufacturer’s protocol. Random primers (Invitrogen) were used for synthesis of complementary DNA. The quantitative real time PCR was performed with a CFX384 Cycler (Bio-Rad Laboratories), using SYBR Green Supermix (Bio-Rad Laboratories) and the following primers: *Pol2* 5’-CTTCCGGGGCCATGTATCTT-3’ 5’-GCTCGATACCCTGCAGGGTCA-3’; *Il15* 5’-GTGACTTTCATCCCAGTTGC-3’ 5’-TCACATTCCTTGCAGCCAGA-3’; *Il15ra* 5’-GAGGTCAGGAAAGAATCCACCT-3’ 5’-AGCAAGGACCATGAAGAGGC-3’; *Il12* 5’-GAGGACTTGAAGATGTACCAG-3’ 5’-TTCTATCTGTGTGAGGAGGGC-3’; *Il18* 5’-CAAACCTTCCAAATCACTTCCT-3’ 5’-TCCTTGAAGTTGACGCAAGA-3’; *Ifng* 5’-GCATTCATGAGTATTGCCAAG-3’ 5’-GGTGGACCACTCGGATGA-3’; *Tgfbr2* 5’-AACGACTTGACCTGTTGCCTGT-3’ 5’-CTTCCGGGGCCATGTATCTT-3’; *Gzmb* 5’-ACACCTCCTTCCTCCCCTTC-3’ 5’-TAGGGACGGGAATGTGGACT-3’; *Tgfb1* 5’-ATGCTAAAGAGGTCACCCGC-3’ 5’-TGCTTCCCGAATGTCTGACG-3’; *Runx3* 5’-TACCTACCACCGAGCCATCA-3’ 5’-TTCTATCTTCTGCCGGTGCC-3’; *Smad7* 5’-TCAAACCAACTGCAGGCTGTC-3’ 5’-TCTTCTCCTCCCAGTATGCCA-3’; *Itga1* 5’-CCACCAAGATGAACGAGCCT-3’ 5’-GGCTGCCCAGCGATATAGAG-3’; *Xcl1* 5’-TGAACTTACAAACCCAGCGG-3’ 5’-TCGCTGCTTTCACCCATTTG-3’; *Cdk6* 5’-TCCTGCTCCAGTCCAGCTAT-3’ 5’-CCACGTCTGAACTTCCACGA-3’; *Cotl1* 5’-ATCACATGGATCGGGGAGGA-3’ 5’-TCCGGTCGCTGATCACAAAT-3’; *Car2* 5’-ACTGGGGATACAGCAAGCAC-3’ 5’-TGCTCTTGGACGCAGCTTTA-3’; *Pmepa1* 5’-TCCTTCATCAGCCGACACAG-3’ 5’-CCACCTGACACCGTACTCTC-3’. Individual samples were run in technical triplicates. Gene expression relative to the housekeeping gene *PolII* was calculated with the ΔCt values as previously described (Pfaffl, 2001).

### In vitro stimulation of NK cells

Liver NK cells (live, CD45+CD3-CD5-Ly6G-CD19-CD115-NK1.1+NKp46+) were sorted with a BD FACSAria™ III sorter and seeded in a 96-well round-bottom plate with 2×10^5^ cells per well. Cells were supplemented with 0.1 ng/ml of IL-15/IL-15R complex recombinant protein (ThermoFischer) in all conditions. For IL-15 stimulation, cells were incubated with 25 ng/ml of IL-15/IL-15R complex and 50 µg/ml of anti-TGF-β blocking antibody (clone 1D11.16.8; BioXCell). For TGF-β stimulation, cells were incubated with 10 ng/ml hTGF-β1 (Peprotech). For the double stimulation, cells were incubated with 25 ng/ml of IL-15/IL-15R complex plus 10 ng/ml hTGF-β1. After 48 h, cells were lysed, and the RNA was collected as described in “Quantitative real-time PCR”.

### Immunofluorescence

Livers were fixed in 4% PFA (Roti-Histofix 4%, Roth) for 4 h and dehydrated in 30% sucrose for 48 h at 4°C. Livers were frozen in Tissue-Tek O.C.T. (Sakura), and 10-µm cryo-sections were cut and dried for 30 min at 37°C. Slides were washed with PBS and blocked/permeabilized with PBS/4% BSA (Roth)/0.1% Triton X-100 RT (Sigma) blocking buffer. Subsequently, slides were washed with PBS/0.05% Tween 20 (Sigma), and stained with goat-anti-mouse-VCAM-1 (R&D, 1:100 in PBS/1% BSA) at 4°C overnight. After incubation, slides were washed and stained with donkey-anti-goat IgG-Alexa Fluor 488 (JacksonImmuno, 1:500 in PBS/1% BSA). Finally, slides were washed and incubated with 0.5 µg/ml 4′,6 diamidine-2-phenylindole (DAPI; Invitrogen) for 5 min, and mounted with ProlongDiamond medium (Invitrogen). Stained slides were scanned using the automated multispectral microscopy system Vectra 3.0 (PerkinElmer). Eight-to-10 representative areas were imaged at 200-fold magnification. Spectral unmixing, cell segmentation and quantification was performed using inForm software (PerkinElmer) as described (Silina et al., 2018).

### Statistics

P-values were calculated using GraphPad Prism software (GraphPad Software Inc.). For comparison of two experimental groups, the unpaired two-tailed Student’s t test was used, for comparison of more than two groups the one-way ANOVA multiple comparison correction was used. P-values smaller than 0.05 were considered significant. * *p* < 0.05; ** *p* < 0.01; *** *p* < 0.001. Data are shown as mean ± SD.

## Notes

### Competing Interest Statement

The authors have declared no competing interest.

