## Supplemental figures for "Conventional NK cells and tissue-resident ILC1s join forces to control liver metastasis"

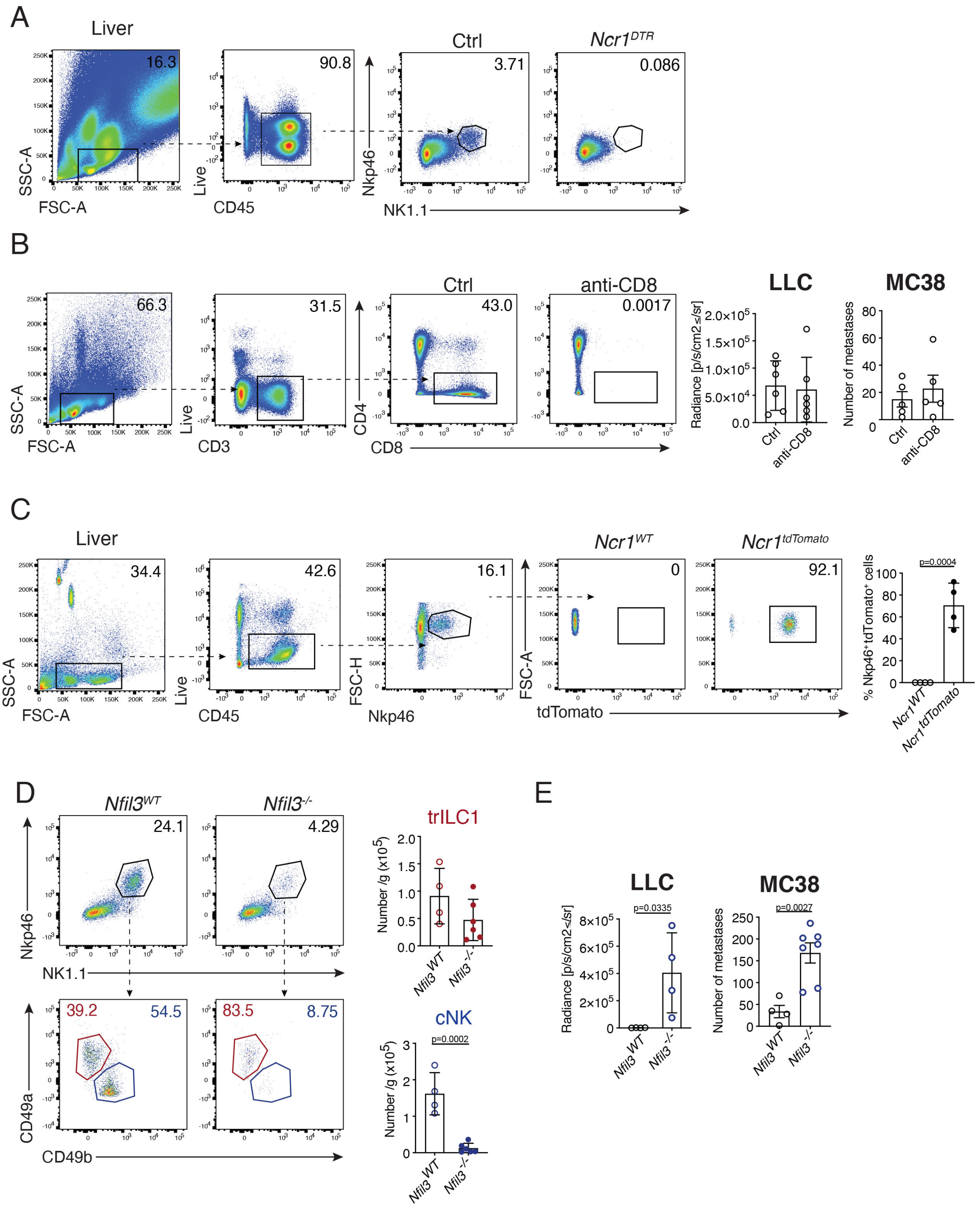

A
Liver
Ctrl
Ncr1DTR
16.3
90.8
3.71
0.086
SSC-A
FSC-A
Live
CD45
Nkp46
NK1.1
B
Ctrl
anti-CD8
66.3
31.5
43.0
0.0017
SSC-A
FSC-A
Live
CD3
CD4
CD8
LLC
MC38
Radiance [p/s/cm2≤/sr]
Number of metastases
C
Liver
Ncr1WT
Ncr1tdTomato
34.4
42.6
16.1
0
92.1
SSC-A
FSC-A
Live
CD45
FSC-H
Nkp46
FSC-A
tdTomato
% Nkp46+tdTomato+ cells
p=0.0004
D
Nfil3WT
Nfil3-/-
24.1
4.29
39.2
54.5
83.5
8.75
Nkp46
NK1.1
CD49a
CD49b
trILC1
cNK
Number /g (x10^5)
p=0.0002
E
LLC
MC38
Radiance [p/s/cm2≤/sr]
Number of metastases
p=0.0335
p=0.0027

**Supplementary Figure 1 (*previous page*). Absence of NK cells, $CD8^+$ T cells, cNKs or trILC1s in different experimental systems, and resulting effect on hepatic metastatic load. Related to Figures 1 and 2.**

**(A)** NK cells were depleted by injection of 250 ng diphtheria toxin intraperitoneally as indicated in Figure 1A. Representative detection of NK cells in livers of control mice ($Ncr1^{iCre/wt}R26R^{wt/wt}$) and $Ncr1^{DTR}$ ($Ncr1^{iiCre/wt}$. $R26R^{iDTR/wt}$) at the endpoint. **(B)** $CD8^+$ T cells were depleted by i.p. injection of 100 μg anti-CD8 (anti-CD8) or isotype control (Ctrl) antibody at -48 h relatively to tumor cell injection. Dot plots: Representative detection of $CD8^+$ T cells depletion in blood of C57BL/6 mice. Bar graphs: Quantification of LLC liver nodules by *ex vivo* IVIS imaging or MC38 macroscopic liver nodules at the endpoint. The bar represents the mean ± SD, symbols represent livers from individual mice, groups consisted of 5-6 mice. One-way analysis of variance (ANOVA), with Tukey's multiple comparisons test. The experiment was performed twice with similar results. **(C)** Representative dot plots and quantification of NK cell in naive livers of $Ncr1^{WT}$ = $Ncr1^{iCre/wt}.R26R^{wt/wt}$; $Ncr1^{tdTomato}$ = $Ncr1^{iCre/wt}.R26R^{Ai14/wt}$ mice. The bar represents the mean ± SD, symbols represent livers from individual mice. Unpaired Student's t-test. Pooled data from two experiments are shown. **(D)** Representative dot plots and quantification of cNKs and trILC1s in naïve livers. Samples were pre-gated on single live $CD45^+lineage^-$ cells and subsequently gated on $NK1.1^+NKp46^+$ cells. cNK = conventional NK cells, $CD49a^-CD49b^+$; trILC1 = tissue-resident ILC1s, $CD49a^+CD49b^-$. The bar represents the mean ± SD, symbols represent livers from individual mice, groups consisted of 4-6 mice. Student's unpaired t test. The experiment was performed twice with similar results. **(E)** Quantification of the LLC- and MC38-derived metastatic burden in livers of $Nfil3^{WT}$ and $Nfil3^{-/-}$ mice as described under (B). The bar represents the mean ± SD, symbols represent livers from individual mice, groups consisted of 4-7 mice. Unpaired Student's t-test. The experiment was performed twice with similar results.

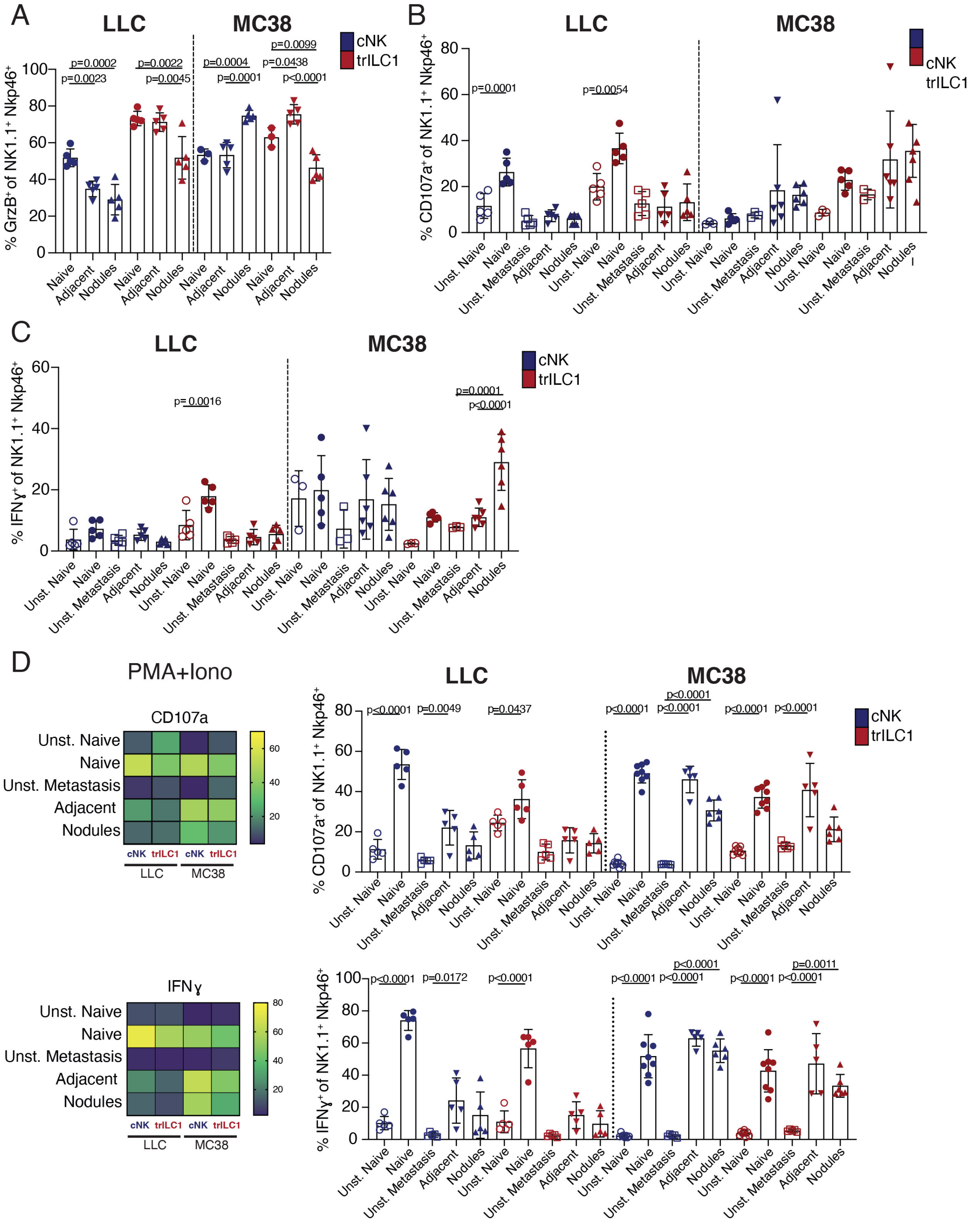

A
LLC
MC38
cNK
trILC1
% GrzB+ of NK1.1+ Nkp46+
p=0.0002
p=0.0023
p=0.0022
p=0.0045
p=0.0004
p=0.0001
p=0.0099
p=0.0438
p<0.0001
Naive
Adjacent
Nodules
B
LLC
MC38
cNK
trILC1
% CD107a+ of NK1.1+ Nkp46+
p=0.0001
p=0.0054
Unst. Naive
Naive
Unst. Metastasis
Adjacent
Nodules
C
LLC
MC38
cNK
trILC1
% IFNγ+ of NK1.1+ Nkp46+
p=0.0016
p=0.0001
p<0.0001
Unst. Naive
Naive
Unst. Metastasis
Adjacent
Nodules
D
PMA+Iono
CD107a
IFNγ
Unst. Naive
Naive
Unst. Metastasis
Adjacent
Nodules
cNK
trILC1
LLC
MC38
LLC
MC38
% CD107a+ of NK1.1+ Nkp46+
p<0.0001
p=0.0049
p=0.0437
p<0.0001
p<0.0001
p<0.0001
p<0.0001
p<0.0001
% IFNγ+ of NK1.1+ Nkp46+
p<0.0001
p=0.0172
p<0.0001
p<0.0001
p<0.0001
p<0.0001
p<0.0001
p<0.0001
p=0.0011

**Supplementary Figure 2 (*previous page*). Effector functions of cNKs and trILC1s in naïve and metastatic livers after stimulation with plate-bound anti-NK1.1 or PMA + ionomycin. Related to Figure 4.**

Naïve and metastatic livers were collected. Metastatic livers were manually dissected to separate the nodules from the adjacent tissue. Tissues were enzymatically processed into a single-cell suspension. Suspensions were stimulated for 4 hours with plate-bound anti-NK1.1 (B, C) or PMA + ionomycin (D), and the expression of effector molecules by cNKs and trILC1s was measured by flow cytometry.
**(A)** Intracellular expression of Granzyme B directly measured directly *ex vivo*. **(B)** Surface expression of CD107a after 4-h stimulation with plate-bound anti-NK1.1. **(C)** Intracellular expression of IFN-γ after 4-h stimulation with plate-bound anti-NK1.1. Unst. = unstimulated. **(D)** Intracellular expression of IFN-γ after 4-h stimulation with 100 ng/ml PMA + 1 $\mu$g/ml ionomycin. Left panel: Heatmap showing the mean proportion of cNKs and trILC1s expressing IFN-γ. Right panel: Quantification of IFN-γ expression by cNKs (blue) and trILC1s (red). Unst. = unstimulated. (A-D) The bar represents the mean ± SD, symbols represent livers from individual mice, groups consisted of 3-8 mice. One-way analysis of variance (ANOVA), with Tukey's multiple comparisons test. The experiment was performed twice with similar results.

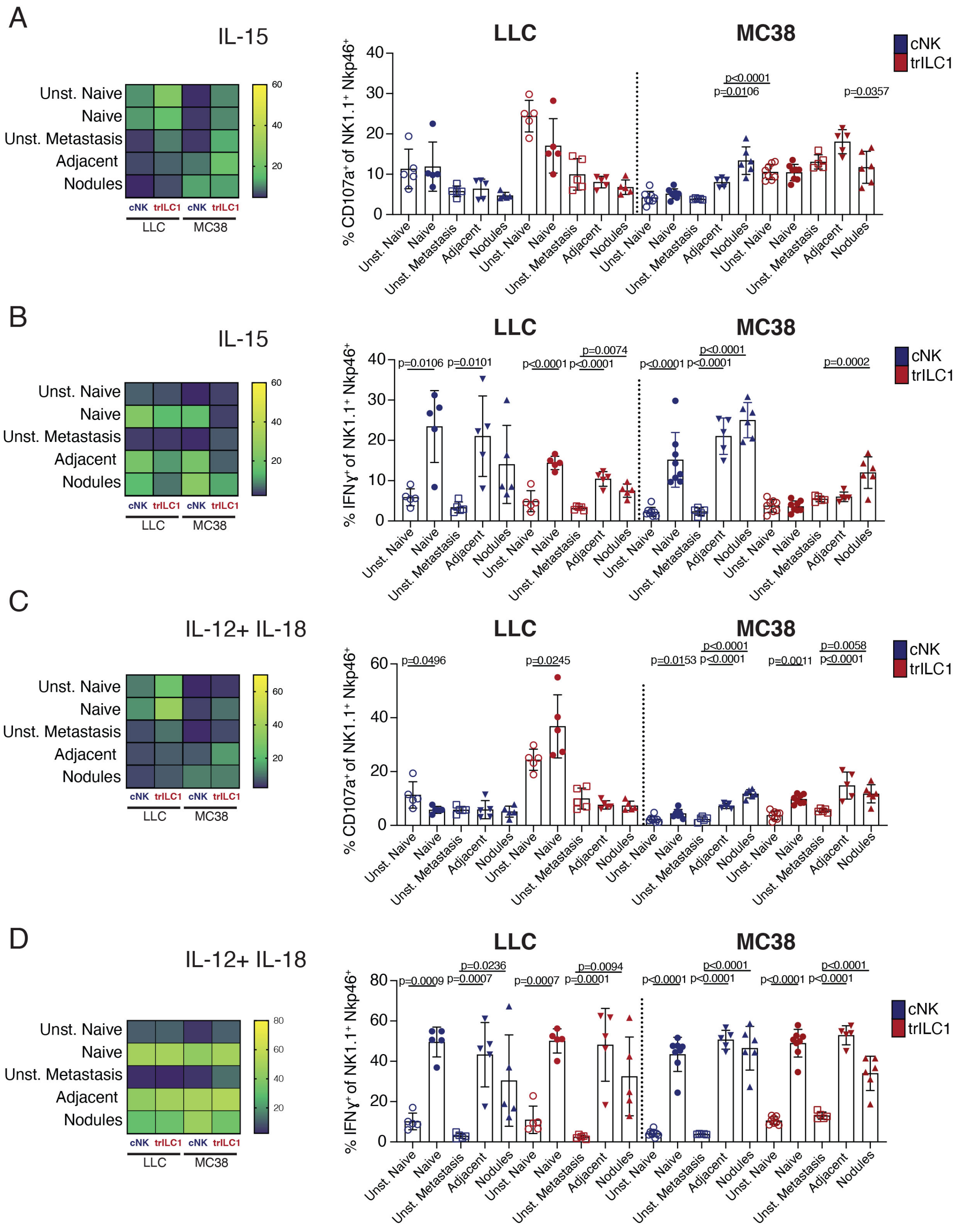
A
IL-15
Unst. Naive
Naive
Unst. Metastasis
Adjacent
Nodules
cNK trILC1 cNK trILC1
LLC MC38
60
40
20
LLC
MC38
cNK
trILC1
% CD107a+ of NK1.1+ Nkp46+
p<0.0001
p=0.0106
p=0.0357
B
IL-15
% IFNγ+ of NK1.1+ Nkp46+
p=0.0106 p=0.0101
p=0.0074
p<0.0001 p<0.0001
p<0.0001
p<0.0001 p<0.0001
p=0.0002
C
IL-12+ IL-18
% CD107a+ of NK1.1+ Nkp46+
p=0.0496
p=0.0245
p=0.0153
p<0.0001 p<0.0001
p=0.0011
p=0.0058 p<0.0001
D
IL-12+ IL-18
% IFNγ+ of NK1.1+ Nkp46+
p=0.0009
p=0.0236 p=0.0007
p=0.0007
p=0.0094 p=0.0001
p<0.0001
p<0.0001 p<0.0001
p<0.0001
p<0.0001 p<0.0001

**Supplementary Figure 3 (*previous page*). Effector functions of cNKs and trILC1s in naïve and metastatic livers after stimulation with IL-15 or IL-12 + IL-18. Related to Figure 4.**

Naïve and metastatic livers were collected. Metastatic livers were manually dissected to separate the nodules from the adjacent tissue. Tissues were enzymatically processed into a single-cell suspension. Suspensions were stimulated for 4 hours with 10 ng/ml IL-15 **(A, B)** or 10 ng/ml IL-12 + 100 ng/ml IL-18 **(C, D)**, and the expression of effector molecules by cNKs and trILC1s was measured by flow cytometry.

**(A)** Surface expression of CD107a after 4-h stimulation with IL-15. **(B)** Intracellular expression of IFN-γ after 4-h stimulation with IL-15. **(C)** Surface expression of CD107a after 4-h stimulation with IL-12 + IL-18. **(D)** Intracellular expression of IFN-γ after 4-h stimulation with IL-12 + Il-18. Unst. = unstimulated. (A-D) Left panels: Heatmap showing the mean proportion of cNKs and trILC1s expressing CD107a or IFN-γ. Right panels: Quantification of CD107a or IFN-γ expression by cNKs (blue) and trILC1s (red). Unst. = unstimulated. The bar represents the mean ± SD, symbols represent livers from individual mice, groups consisted of 5-8 mice. One-way analysis of variance (ANOVA), with Tukey's multiple comparisons test. The experiment was performed once.

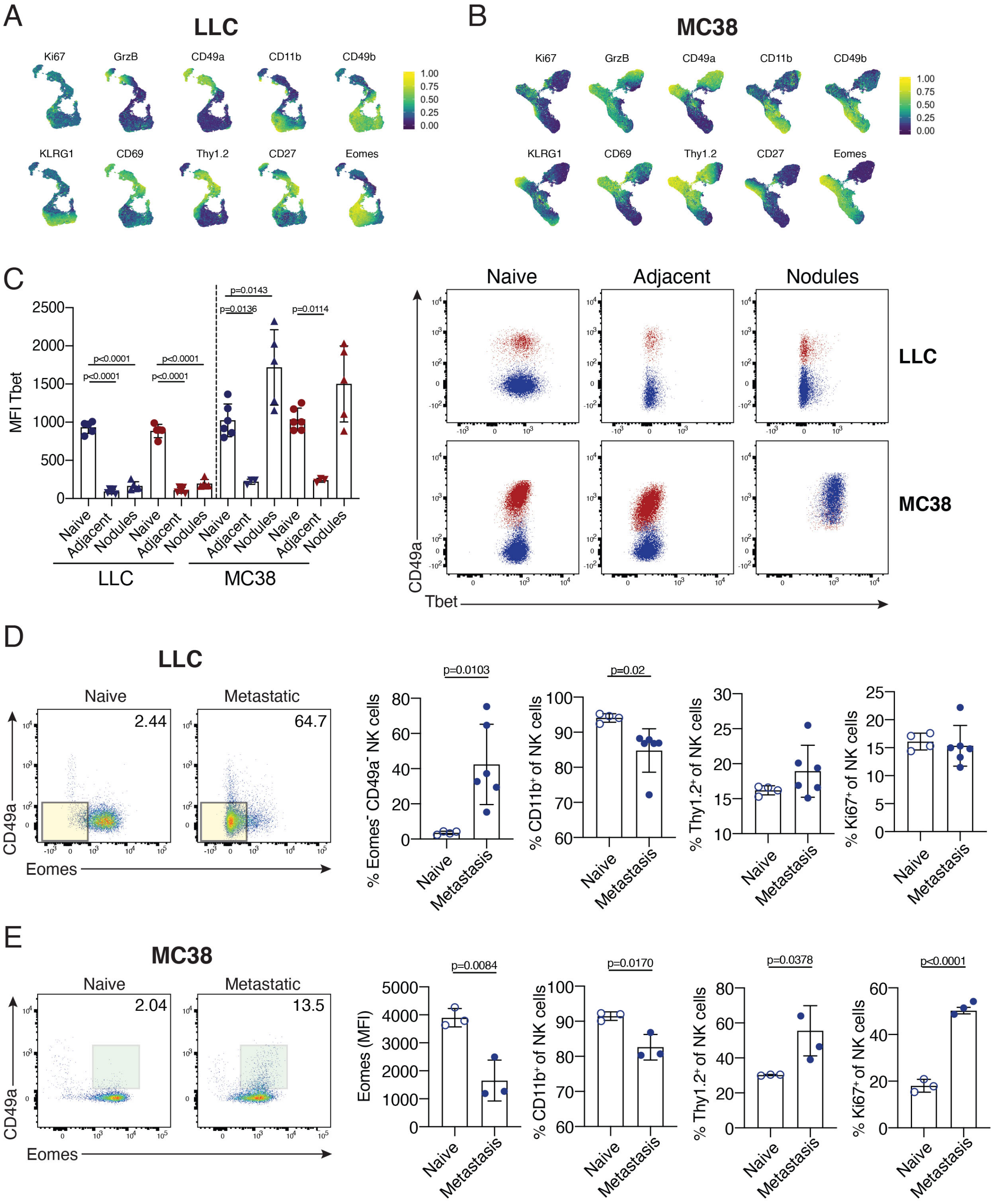

A
LLC
Ki67
GrzB
CD49a
CD11b
CD49b
KLRG1
CD69
Thy1.2
CD27
Eomes
1.00
0.75
0.50
0.25
0.00
B
MC38
C
MFI Tbet
p<0.0001
p=0.0143
p=0.0136
p=0.0114
Naive
Adjacent
Nodules
LLC
MC38
CD49a
Tbet
D
LLC
Naive
Metastatic
2.44
64.7
Eomes
p=0.0103
p=0.02
% Eomes- CD49a- NK cells
% CD11b+ of NK cells
% Thy1.2+ of NK cells
% Ki67+ of NK cells
Metastasis
E
MC38
2.04
13.5
p=0.0084
p=0.0170
p=0.0378
p<0.0001
Eomes (MFI)

**Supplementary Figure 4 (*previous page*). Multi-parameter single-cell mapping of cNKs and trILC1s cells in blood and livers from naïve and metastatic mice. Related to Figure 5.**

Naïve and metastatic livers were collected. Metastatic livers (day 21) were manually dissected to separate the nodules from the adjacent tissue, and tissues were enzymatically processed into a single-cell suspension. NK cells were analyzed by multi-parameter single-cell mapping using flow cytometry. UMAP visualization of markers after gating on single, live, $CD45^{+}lin^{-}NK1.1^{+}NKp46^{+}$cells. **(A)** LLC-metastatic livers. **(B)** MC38-metastatic livers. **(C)** Expression of Tbet by cNKs (blue) and trILC1s (red) in metastatic livers. The bar represents the mean ± SD, symbols represent livers from individual mice, groups consisted of 3-5 mice. One-way analysis of variance (ANOVA), with Tukey's multiple comparisons test. The experiment was performed twice with similar results. **(D)** Flow cytometry analysis of $CD49a^{-}Eomes^{-}$ population from blood of mice with naïve or LLC-metastatic livers. Left panels: Highlighted in yellow is the $CD49a^{-}Eomes^{-}$ population. Right panels: Quantification of $CD49a^{-}Eomes^{-}$ population, $CD11b^{+}$, $Thy1.2^{+}$ and $Ki67^{+}$ cells in cNKs (blue). The bar represents the mean ± SD, symbols represent livers from individual mice, groups consisted of 4-5 mice. Unpaired Student's t-test. The experiment was performed twice with similar results. **(E)** Flow cytometry analysis of $CD49a^{+}Eomes^{+}$ population from blood of mice with naïve or MC38-metastatic livers. Left panels: Highlighted in green is the $CD49a^{+}Eomes^{+}$ population. Right panels: Quantification of Eomes expression, $CD11b^{+}$, $Thy1.2^{+}$ and $Ki67^{+}$ cells in cNKs (blue). The bar represents the mean ± SD, symbols represent livers from individual mice, groups consisted of 3 mice. Unpaired Student's t-test. The experiment was performed twice with similar results.

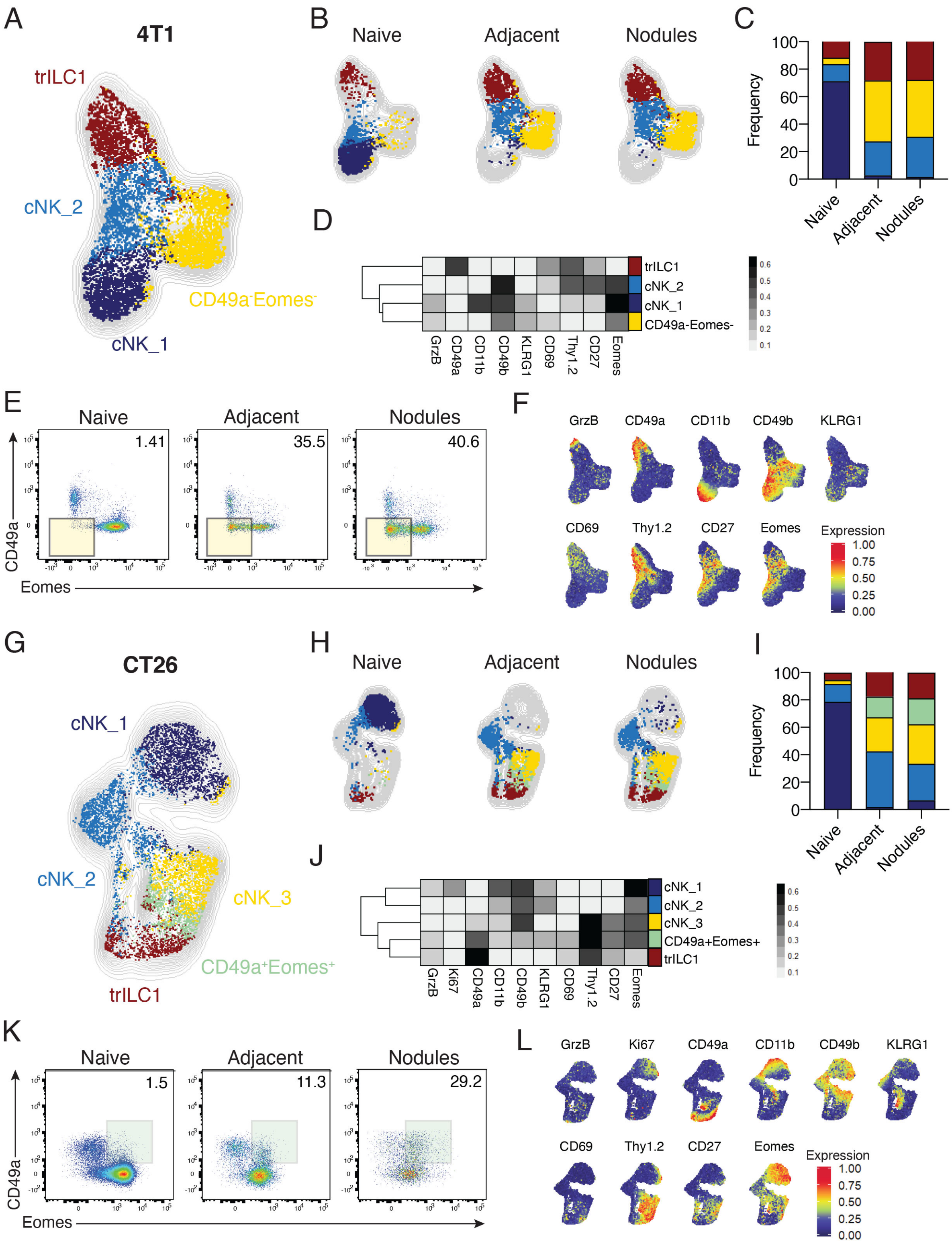
A
4T1
trILC1
cNK_2
CD49a-Eomes-
cNK_1
B
Naive
Adjacent
Nodules
C
Frequency
Naive
Adjacent
Nodules
D
trILC1
cNK_2
cNK_1
CD49a-Eomes-
GrzB
CD49a
CD11b
CD49b
KLRG1
CD69
Thy1.2
CD27
Eomes
E
Naive
Adjacent
Nodules
1.41
35.5
40.6
CD49a
Eomes
F
GrzB
CD49a
CD11b
CD49b
KLRG1
CD69
Thy1.2
CD27
Eomes
Expression
G
CT26
cNK_1
cNK_2
cNK_3
CD49a+Eomes+
trILC1
H
Naive
Adjacent
Nodules
I
Frequency
Naive
Adjacent
Nodules
J
cNK_1
cNK_2
cNK_3
CD49a+Eomes+
trILC1
GrzB
Ki67
CD49a
CD11b
CD49b
KLRG1
CD69
Thy1.2
CD27
Eomes
K
Naive
Adjacent
Nodules
1.5
11.3
29.2
CD49a
Eomes
L
GrzB
Ki67
CD49a
CD11b
CD49b
KLRG1
CD69
Thy1.2
CD27
Eomes
Expression

M

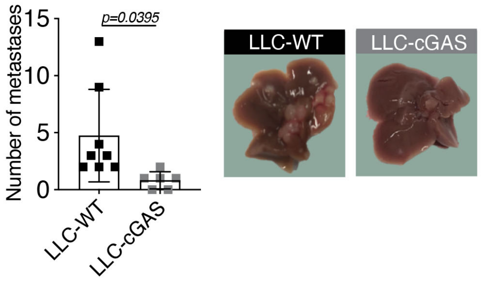


N

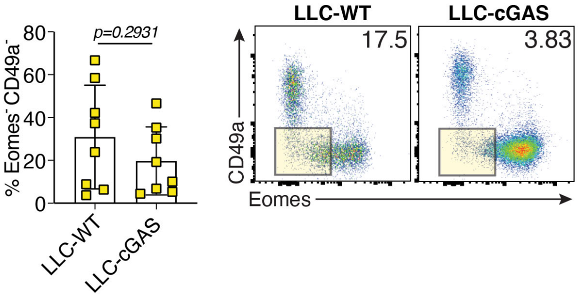


**Supplementary Figure 5. Metastasis drives the emergence of unique cNK cell populations. Related to Figure 5.**

Naïve and metastatic livers were collected. Metastatic livers (day 21) were manually dissected to separate the nodules from the adjacent tissue, and tissues were enzymatically processed into a single-cell suspension. NK cells were analyzed by multi-parameter single-cell mapping using flow cytometry. Samples were pre-gated on single live CD45$^{+}$lineage$^{-}$ cells and subsequently gated on NK1.1$^{+}$NKp46$^{+}$ cells. cNK = conventional NK cells, CD49a$^{-}$CD49b$^{+}$ ; trILC1 = tissue-resident ILC1s, CD49a$^{+}$CD49b$^{-}$. **(A-F)** 4T1-metastatic and control livers. **(G-L)** CT26-metastatic and control livers. **(A, G)** UMAP maps overlaid with FlowSOM-guided manual metaclusters displaying cNKs and trILC1s from all samples. **(B, H)** UMAP maps overlaid with FlowSOM-guided manual metaclusters separated by sample category (naïve, adjacent, nodules). **(C, I)** Relative frequency of each cluster in the different sample categories (naïve, adjacent, nodules). **(D, J)** Heatmap summary of median marker expression values of the different markers analyzed for each cluster. **(E)** Highlighted in yellow is the CD49a$^{-}$Eomes$^{-}$ population observed in 4T1 adjacent tissue and nodules. **(F)** UMAP visualization of markers after gating on single, live, CD45$^{+}$lin$^{-}$NK1.1$^{+}$NKp46$^{+}$cells. **(K)** Highlighted in green is the CD49a$^{+}$Eomes$^{+}$ population observed in CT26 nodules. **(L)** UMAP visualization of markers after gating on single, live, CD45$^{+}$lin$^{-}$NK1.1$^{+}$NKp46$^{+}$cells in mice. Experimental groups consisted of 4-5 mice. The experiment was performed twice with similar results. **(M)** Left panel: Macroscopic quantification of LLC-WT and LLC-cGAS metastatic nodules in the liver of C57BL/6 mice 21 days after tumor cell injection. Bars show the mean ± SD. Each symbol represents an individual mouse. One-way analysis of variance (ANOVA), with Tukey's multiple comparisons test. The experiment was performed twice with similar results. Right panels: Representative images of metastatic livers from each group at the endpoint. LLC-WT = control LLC cells; LLC-cGAS = LLC cells overexpressing cGAS (*Mb21d1*). **(N)** Left panel: Percentage of CD49a$^{-}$Eomes$^{-}$ NK cells in LLC-WT and LLC-cGAS nodules. Bars show the mean ± SD. Each symbol represents an individual mouse. One-way analysis of variance (ANOVA), with Tukey's multiple comparisons test. The experiment was performed twice with similar results. Right panel: Representative dot plots of cNKs and trILC1s in LLC-WT and LLC-cGAS nodules, with the CD49a$^{-}$Eomes$^{-}$ population highlighted in yellow.

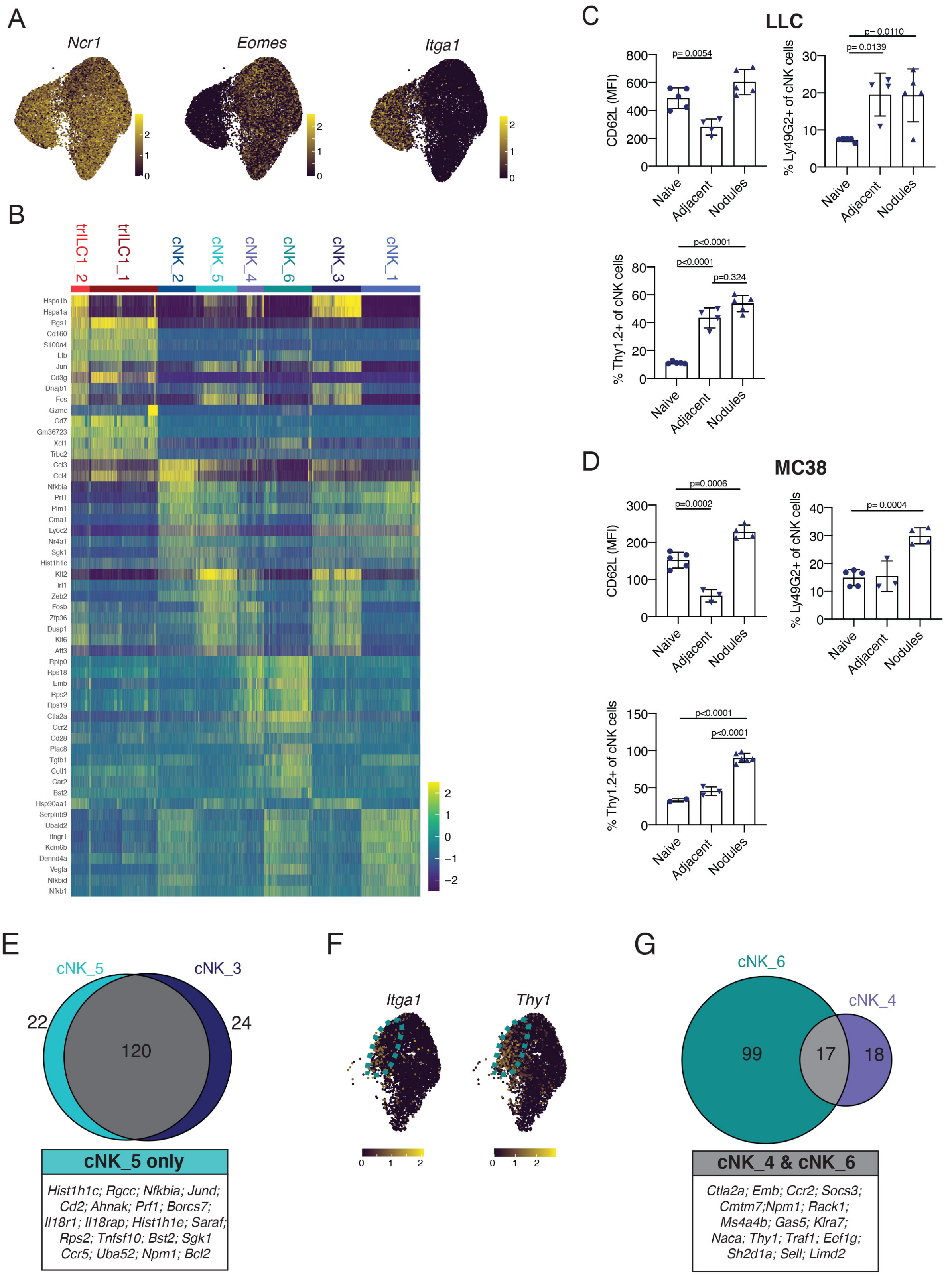
A
Ncr1
Eomes
Itga1
B
trILC1_2
trILC1_1
cNK_2
cNK_5
cNK_4
cNK_6
cNK_3
cNK_1
Hspa1b
Hspa1a
Rgs1
Cd160
S100a4
Ltb
Jun
Cd3g
Dnajb1
Fos
Gzmc
Cd7
Gm36723
Xcl1
Trbc2
Ccl3
Ccl4
Nfkbia
Prf1
Pim1
Cma1
Ly6c2
Nr4a1
Sgk1
Hist1h1c
Klf2
Irf1
Zeb2
Fosb
Zfp36
Dusp1
Klf6
Atf3
Rplp0
Rps18
Emb
Rps2
Rps19
Ctla2a
Ccr2
Cd28
Plac8
Tgfb1
Cotl1
Car2
Bst2
Hsp90aa1
Serpinb9
Ubald2
Ifngr1
Kdm6b
Dennd4a
Vegfa
Nfkbid
Nfkb1
C
LLC
CD62L (MFI)
p= 0.0054
% Ly49G2+ of cNK cells
p= 0.0110
p= 0.0139
% Thy1.2+ of cNK cells
p<0.0001
p<0.0001
p=0.324
Naive
Adjacent
Nodules
D
MC38
CD62L (MFI)
p=0.0006
p=0.0002
% Ly49G2+ of cNK cells
p= 0.0004
% Thy1.2+ of cNK cells
p<0.0001
p<0.0001
Naive
Adjacent
Nodules
E
cNK_5
cNK_3
22
120
24
cNK_5 only
Hist1h1c; Rgcc; Nfkbia; Jund; Cd2; Ahnak; Prf1; Borcs7; Il18r1; Il18rap; Hist1h1e; Saraf; Rps2; Tnfsf10; Bst2; Sgk1 Ccr5; Uba52; Npm1; Bcl2
F
Itga1
Thy1
G
cNK_6
cNK_4
99
17
18
cNK_4 & cNK_6
Ctla2a; Emb; Ccr2; Socs3; Cmtm7;Npm1; Rack1; Ms4a4b; Gas5; Klra7; Naca; Thy1; Traf1; Eef1g; Sh2d1a; Sell; Limd2

**Supplementary Figure 6 (*previous page*). Single-cell RNA-sequencing reveals novel transcriptional signatures of cNKs in the hepatic metastatic niche. Related to Figure 6.**

Single-cell RNA-sequencing was performed on NK cells sorted from naïve, LLC-metastatic and MC38-metastatic livers (6 mice per condition, pooled into 1 sample for droplet encapsulation and library preparation). **(A)** UMAP analysis of all NK cells with overlaid expression of key lineage markers: Nkp46 (*Ncr1*), CD49a (*Itga1*) and Eomes (*Eomes*). **(B)** Heatmap showing the expression of the top-10 genes for each cluster. Cells are plotted in columns and grouped by cluster, and the genes are plotted in rows. **(C, D)** Naïve and metastatic livers were collected. Metastatic livers (day 21) were manually dissected to separate the nodules from the adjacent tissue, and tissues were enzymatically processed into a single-cell suspension. NK cells were analyzed by flow cytometry. Expression of $CD62L^{+}$, $Thy1.2^{+}$ and $Ly49G2^{+}$ by hepatic cNKs. **(C)** LLC-metastatic liver. **(D)** MC38-metastatic liver. The bar represents the mean ± SD, symbols represent livers from individual mice, groups consisted of 2-6 mice. One-way analysis of variance (ANOVA), with Tukey's multiple comparisons test. The experiment was performed twice with similar results. **(E)** Venn diagram showing differentially expressed genes (DEG) in clusters cNK_3 and cNK_5. The box shows the list of genes exclusive to cNK_5. **(F)** UMAP analysis of cNKs from MC38-metastatic livers with overlaid expression of CD49a (*Itga1*) and Thy1.2 (*Thy1*). The dotted line marks cluster cNK_6. **(G)** Venn diagram showing the differentially expressed genes in clusters cNK_4 and cNK_6. The box shows the list of genes shared by both clusters.

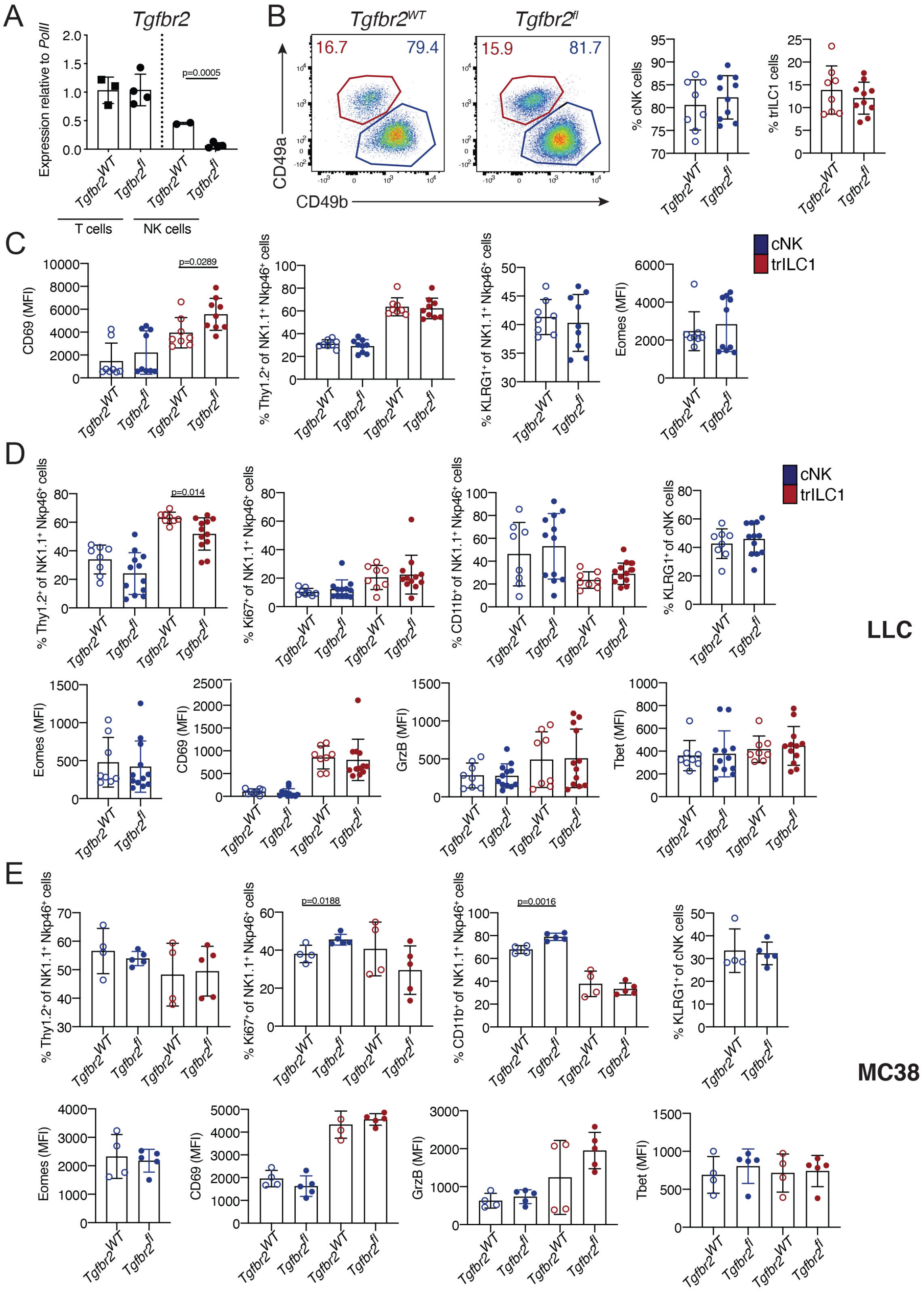
A
Tgfbr2
Expression relative to PolII
2.0
1.5
1.0
0.5
0.0
p=0.0005
Tgfbr2WT
Tgfbr2fl
T cells
NK cells
B
Tgfbr2WT
Tgfbr2fl
16.7
79.4
15.9
81.7
CD49a
CD49b
% cNK cells
% trILC1 cells
C
CD69 (MFI)
p=0.0289
% Thy1.2+ of NK1.1+ Nkp46+ cells
% KLRG1+ of NK1.1+ Nkp46+ cells
Eomes (MFI)
cNK
trILC1
D
% Thy1.2+ of NK1.1+ Nkp46+ cells
p=0.014
% Ki67+ of NK1.1+ Nkp46+ cells
% CD11b+ of NK1.1+ Nkp46+ cells
% KLRG1+ of cNK cells
cNK
trILC1
LLC
Eomes (MFI)
CD69 (MFI)
GrzB (MFI)
Tbet (MFI)
E
% Thy1.2+ of NK1.1+ Nkp46+ cells
% Ki67+ of NK1.1+ Nkp46+ cells
p=0.0188
% CD11b+ of NK1.1+ Nkp46+ cells
p=0.0016
% KLRG1+ of cNK cells
MC38
Eomes (MFI)
CD69 (MFI)
GrzB (MFI)
Tbet (MFI)

**Supplementary Figure 7 (*previous page*). TGF-β modulates the fate of CD49a$^+$Eomes$^+$ cNKs in metastatic nodules. Related to Figure 7.**

**(A)** Expression of *Tgfbr2* transcripts by T and NK cells sorted from naïve livers. *$Tgfbr2^{WT}$* = *$Ncr1^{iCre/wt}$.$Tgfbr2^{wt/wt}$*; *$Tgfbr2^{fl}$* = *$Ncr1^{iCre/wt}$.$Tgfbr2^{fl/fl}$*. **(B)** Representative dot plots and quantification of cNKs and trILC1s in naïve livers of *$Tgfbr2^{WT}$* and *$Tgfbr2^{fl}$* mice. Samples were gated on single, live, CD45$^+$lin$^-$NK1.1$^+$NKp46$^+$cells. cNK = conventional NK cells, CD49a$^-$CD49b$^+$; trILC1 = tissue-resident ILC1s, CD49a$^+$CD49b$^-$. The bar represents the mean ± SD, symbols represent livers from individual mice, groups consisted of 8-10 mice. Unpaired Student's t-test. **(C)** Expressing CD69$^+$, Thy1.2$^+$, KLRG1$^+$ cells and Eomes by cNKs (blue) and trILC1s (red) from naïve livers. *$Tgfbr2^{WT}$* = *$Ncr1^{iCre/wt}$.$Tgfbr2^{wt/wt}$*; *$Tgfbr2^{fl}$* = *$Ncr1^{iCre/wt}$.$Tgfbr2^{fl/fl}$*. The bar represents the mean ± SD, symbols represent livers from individual mice, groups consisted of 8-9 mice. One-way analysis of variance (ANOVA), with Tukey's multiple comparisons test. **(D-E)** Upper panels: Proportion of cells expressing Thy1.2$^+$, Ki67$^+$, CD11b$^+$ or KLRG1$^+$. Lower panels: Expression of Eomes, CD69, GrzB or Tbet expression in cNKs (blue) and trILC1s (red) from metastatic livers. *$Tgfbr2^{WT}$* = *$Ncr1^{iCre/wt}$.$Tgfbr2^{wt/wt}$*; *$Tgfbr2^{fl}$* = *$Ncr1^{iCre/wt}$.$Tgfbr2^{fl/fl}$*. **(D)** LLC-metastatic livers. **(E)** MC38-metastatic livers. The bar represents the mean ± SD, symbols represent livers from individual mice, groups consisted of 5-12 mice. One-way analysis of variance (ANOVA), with Tukey's multiple comparisons test.
