## Supplemental table for "Conventional NK cells and tissue-resident ILC1s join forces to control liver metastasis"

| trILC1_1 | | |
|---|---|---|
| gene | p_val | avg_logFC |
| Cd3g | 0 | 1.6244 |
| Rgs1 | 0 | 1.4893 |
| Gzmc | 0 | 1.0602 |
| S100a4 | 0 | 1.0595 |
| Cd160 | 0 | 1.0065 |
| Cd7 | 0 | 0.9202 |
| Gm36723 | 0 | 0.8701 |
| Xcl1 | 0 | 0.8672 |
| Ltb | 0 | 0.8355 |
| Trbc2 | 0 | 0.8266 |
| Osgin1 | 0 | 0.7474 |
| Gzmb | 0 | 0.7443 |
| Itga1 | 0 | 0.7070 |
| Ly6e | 0 | 0.6577 |
| Il21r | 0 | 0.6394 |
| Ckb | 0 | 0.5931 |
| Rgs2 | 0 | 0.5789 |
| Trbc1 | 0 | 0.5779 |
| Nedd9 | 0 | 0.5756 |
| Cd226 | 0 | 0.5738 |
| Klrc1 | 0 | 0.5694 |
| Ccl4 | 1.3E-257 | 0.5503 |
| S100a6 | 0 | 0.5479 |
| Cxcr3 | 0 | 0.5165 |
| Stk17b | 0 | 0.5001 |
| Irf1 | 5.4E-209 | -0.5458 |
| Ctla2a | 9.4E-162 | -0.5505 |
| Eomes | 0 | -0.5524 |
| Cd2 | 0 | -0.5751 |
| Anxa2 | 0 | -0.5789 |
| Sell | 0 | -0.5861 |
| Emp3 | 0 | -0.5972 |
| Ms4a4b | 0 | -0.6070 |
| Vim | 0 | -0.6409 |
| Itga4 | 0 | -0.7057 |
| Lgals1 | 0 | -0.7561 |
| Klra13-ps | 0 | -0.7580 |
| Sgk1 | 0 | -0.7668 |
| Klra7 | 0 | -0.7740 |
| Zeb2 | 0 | -0.8126 |
| Cma1 | 0 | -0.8313 |
| Hspa1b | 6.5E-69 | -0.9003 |
| Ly6c2 | 0 | -0.9357 |
| Nfkbia | 0 | -0.9376 |
| Hspa1a | 4.8E-94 | -1.0335 |
| Klra8 | 0 | -1.0413 |
| Prf1 | 0 | -1.1552 |
| Klra4 | 0 | -1.3735 |
| Klf2 | 0 | -1.5987 |

| trILC1_2 | | |
|---|---|---|
| gene | p_val | avg_logFC |
| Hspa1b | 0 | 1.3944 |
| Hspa1a | 0 | 1.2469 |
| Rgs1 | 0 | 1.0219 |
| Dnajb1 | 2.1E-257 | 1.0105 |
| Jun | 1.0E-277 | 0.9694 |
| Cd160 | 0 | 0.9236 |
| Cd3g | 6.7E-267 | 0.8895 |
| Fos | 2.7E-215 | 0.8593 |
| S100a4 | 0 | 0.7820 |
| Ltb | 0 | 0.7586 |
| Cd7 | 0 | 0.7302 |
| Gm36723 | 0 | 0.7164 |
| Rgs2 | 4.1E-160 | 0.6949 |
| Cited4 | 3.2E-243 | 0.6761 |
| Trbc2 | 1.3E-188 | 0.6401 |
| Cxcr3 | 0 | 0.6230 |
| Dnaja1 | 6.6E-113 | 0.5712 |
| Hsph1 | 1.8E-196 | 0.5519 |
| Cd226 | 0 | 0.5454 |
| Ly6e | 9.9E-229 | 0.5333 |
| Hsp90aa1 | 3.4E-136 | 0.5254 |
| Klf6 | 5.9E-148 | 0.5235 |
| Klrc1 | 1.0E-148 | 0.5086 |
| Dusp1 | 7.1E-93 | 0.5074 |
| Ckb | 0 | 0.5072 |
| Emp3 | 7.6E-113 | -0.4676 |
| Vim | 1.6E-89 | -0.4709 |
| Ctla2a | 1.0E-53 | -0.4902 |
| Sell | 4.5E-139 | -0.5024 |
| Anxa2 | 1.2E-171 | -0.5237 |
| Pim1 | 7.0E-144 | -0.5822 |
| Itga4 | 1.8E-148 | -0.5974 |
| Kdm6b | 8.9E-132 | -0.6119 |
| Serpinb9 | 2.8E-126 | -0.6149 |
| Sgk1 | 4.4E-90 | -0.6165 |
| Klf2 | 1.8E-47 | -0.6212 |
| Klra13-ps | 1.2E-121 | -0.6315 |
| Gzma | 6.1E-148 | -0.6332 |
| Klra7 | 5.1E-91 | -0.6437 |
| Ubald2 | 1.1E-156 | -0.6485 |
| Zeb2 | 5.8E-151 | -0.6502 |
| Lgals1 | 2.6E-192 | -0.6955 |
| Cma1 | 5.8E-148 | -0.7088 |
| Ifngr1 | 3.3E-254 | -0.7537 |
| Ly6c2 | 1.5E-77 | -0.7759 |
| Klra8 | 7.9E-122 | -0.8485 |
| Nfkbia | 5.5E-252 | -0.9858 |
| Prf1 | 2.9E-297 | -1.1014 |
| Klra4 | 5.6E-175 | -1.1675 |

| cNK_1 | | |
|---|---|---|
| gene | p_val | avg_logFC |
| Prf1 | 0 | 0.6108 |
| Serpinb9 | 0 | 0.6096 |
| Ubald2 | 0 | 0.5878 |
| Ifngr1 | 0 | 0.5402 |
| Kdm6b | 0 | 0.5225 |
| Dennd4a | 0 | 0.5012 |
| Vegfa | 0 | 0.4724 |
| Nfkbid | 2.6E-292 | 0.4300 |
| Nfkb1 | 3.2E-246 | 0.3860 |
| Ly6c2 | 1.2E-146 | 0.3750 |
| Id2 | 8.1E-251 | 0.3707 |
| Gzma | 1.8E-296 | 0.3673 |
| Serpinb6b | 3.9E-152 | 0.3672 |
| Nfkbia | 2.4E-254 | 0.3482 |
| Bcl2a1b | 1.8E-101 | 0.3477 |
| Dusp2 | 9.0E-178 | 0.3445 |
| Orai1 | 1.5E-183 | 0.3442 |
| Btg1 | 3.7E-288 | 0.3373 |
| Ccl5 | 1.5E-276 | 0.3328 |
| Sh2d2a | 6.1E-210 | 0.3306 |
| Rap1b | 1.2E-270 | 0.3256 |
| Cma1 | 1.0E-215 | 0.3180 |
| Irf8 | 0 | 0.3146 |
| Cd9 | 5.3E-189 | 0.3116 |
| Ppig | 3.5E-133 | 0.3084 |
| S100a4 | 1.9E-114 | -0.5067 |
| Dusp1 | 5.9E-149 | -0.5192 |
| Btg2 | 0 | -0.5248 |
| Ccl4 | 3.4E-30 | -0.5383 |
| Xcl1 | 1.2E-163 | -0.5419 |
| Zfp36 | 2.0E-301 | -0.5741 |
| Ly6e | 0 | -0.5769 |
| Cd69 | 5.9E-281 | -0.6065 |
| Klf2 | 1.3E-41 | -0.6281 |
| Cd160 | 6.0E-254 | -0.6446 |
| Ifng | 1.4E-154 | -0.6457 |
| Ccl3 | 1.5E-44 | -0.7037 |
| Klf6 | 0 | -0.7241 |
| Ltb | 0 | -0.7453 |
| Dnajb1 | 9.4E-183 | -0.7537 |
| Atf3 | 1.5E-170 | -0.7668 |
| Fosb | 0 | -0.8091 |
| Gzmc | 9.4E-103 | -0.8812 |
| Cd3g | 5.9E-169 | -1.0074 |
| Rgs1 | 3.7E-220 | -1.0897 |
| Jun | 0 | -1.4354 |
| Fos | 0 | -1.4808 |
| Hspa1a | 8.9E-266 | -1.6612 |
| Hspa1b | 0 | -1.9922 |

| cNK_2 | | |
|---|---|---|
| gene | p_val | avg_logFC |
| Ccl3 | 0 | 1.1094 |
| Ccl4 | 0 | 1.0389 |
| Nfkbia | 0 | 0.6404 |
| Prf1 | 0 | 0.5645 |
| Ly6c2 | 1.5E-161 | 0.4780 |
| Sgk1 | 1.1E-156 | 0.4383 |
| Cma1 | 5.7E-209 | 0.4364 |
| Nr4a1 | 3.5E-161 | 0.4173 |
| Pim1 | 1.2E-252 | 0.4168 |
| Hist1h1c | 5.7E-81 | 0.3780 |
| Klra9 | 2.3E-100 | 0.3776 |
| Dusp2 | 1.2E-95 | 0.3748 |
| Icam1 | 1.4E-117 | 0.3633 |
| Gzma | 1.7E-207 | 0.3557 |
| Nfkbid | 8.0E-118 | 0.3517 |
| Lgals1 | 5.8E-162 | 0.3265 |
| Gadd45b | 4.3E-44 | 0.3157 |
| Emp3 | 6.7E-170 | 0.3030 |
| Nfkbiz | 3.0E-98 | 0.3029 |
| Ubald2 | 3.6E-122 | 0.2946 |
| Bhlhe40 | 1.9E-131 | 0.2915 |
| Klrg1 | 1.1E-138 | 0.2792 |
| Serpinb9 | 2.4E-90 | 0.2699 |
| Klra8 | 1.5E-49 | 0.2653 |
| Rap1b | 2.5E-122 | 0.2569 |
| Cxcr3 | 3.2E-138 | -0.3255 |
| Serpina3g | 6.9E-96 | -0.3486 |
| Klf6 | 2.3E-63 | -0.3626 |
| Trbc1 | 4.6E-123 | -0.3770 |
| Rgs2 | 1.3E-74 | -0.3880 |
| Xcl1 | 5.9E-77 | -0.4070 |
| Gm36723 | 1.6E-136 | -0.4309 |
| Ctla2a | 4.5E-64 | -0.4401 |
| Emb | 1.0E-102 | -0.4489 |
| Trbc2 | 6.4E-92 | -0.4890 |
| Cd7 | 1.7E-165 | -0.5035 |
| S100a4 | 1.2E-85 | -0.5073 |
| Fosb | 7.3E-121 | -0.5273 |
| Dnajb1 | 1.2E-79 | -0.5483 |
| Ly6e | 4.2E-228 | -0.5667 |
| Cd160 | 1.5E-155 | -0.6115 |
| Ltb | 2.9E-233 | -0.6992 |
| Jun | 1.9E-107 | -0.7451 |
| Gzmc | 1.8E-58 | -0.8379 |
| Cd3g | 1.8E-107 | -0.9705 |
| Rgs1 | 2.2E-153 | -1.0460 |
| Fos | 1.1E-170 | -1.0618 |
| Hspa1a | 9.0E-129 | -1.4165 |
| Hspa1b | 8.2E-167 | -1.5368 |

| cNK_3 | | |
|---|---|---|
| gene | p_val | avg_logFC |
| Hspa1a | 0 | 1.9683 |
| Hspa1b | 0 | 1.7944 |
| Klf2 | 0 | 0.8897 |
| Fos | 0 | 0.8117 |
| Dnajb1 | 0 | 0.7946 |
| Jun | 0 | 0.7026 |
| Atf3 | 2.7E-210 | 0.6629 |
| Zeb2 | 0 | 0.5890 |
| Hsp90aa1 | 0 | 0.5376 |
| Klf6 | 0 | 0.5173 |
| Dusp1 | 2.8E-277 | 0.4947 |
| Dnaja1 | 1.9E-196 | 0.4771 |
| Fosb | 5.8E-203 | 0.4648 |
| Irf1 | 7.2E-205 | 0.4509 |
| Ifng | 4.1E-96 | 0.4436 |
| Hspa8 | 1.8E-269 | 0.4392 |
| Ly6c2 | 1.2E-162 | 0.4166 |
| Zfp36 | 5.3E-186 | 0.4040 |
| Rhob | 4.6E-166 | 0.3793 |
| Klra4 | 2.8E-128 | 0.3755 |
| Ubc | 3.5E-215 | 0.3644 |
| Hsp90ab1 | 2.8E-258 | 0.3522 |
| Cma1 | 1.9E-201 | 0.3360 |
| Egr1 | 4.1E-83 | 0.3295 |
| Ier5 | 5.1E-170 | 0.3184 |
| Osgin1 | 1.2E-162 | -0.3622 |
| Srgn | 1.6E-268 | -0.3671 |
| Serpinb9 | 9.8E-99 | -0.3673 |
| Ccl4 | 9.5E-43 | -0.3674 |
| Litaf | 4.1E-206 | -0.3679 |
| Tgfb1 | 5.3E-141 | -0.3726 |
| Neurl3 | 7.5E-139 | -0.3745 |
| Smad7 | 9.3E-153 | -0.3848 |
| Dusp5 | 5.4E-207 | -0.3990 |
| Trbc1 | 6.2E-171 | -0.3998 |
| Dennd4a | 1.2E-204 | -0.4063 |
| Bcl2a1b | 6.2E-116 | -0.4329 |
| Trbc2 | 7.5E-87 | -0.4436 |
| Cd7 | 2.8E-146 | -0.4456 |
| Gm36723 | 3.7E-179 | -0.4477 |
| Prdx6 | 1.2E-266 | -0.4920 |
| Ly6e | 6.3E-229 | -0.5094 |
| S100a4 | 8.6E-114 | -0.5329 |
| Cd160 | 5.1E-158 | -0.5788 |
| Ltb | 3.7E-261 | -0.6604 |
| Xcl1 | 2.6E-206 | -0.6623 |
| Rgs1 | 3.0E-60 | -0.6642 |
| Gzmc | 1.0E-87 | -0.8657 |
| Cd3g | 6.9E-135 | -0.9767 |

| cNK_4 | | |
|---|---|---|
| gene | p_val | avg_logFC |
| Emb | 1.4E-250 | 0.5349 |
| Fosb | 7.4E-179 | 0.5275 |
| Ctla2a | 2.0E-170 | 0.5273 |
| Fos | 8.8E-157 | 0.4433 |
| Cd28 | 1.5E-141 | 0.4272 |
| Ccr2 | 3.2E-152 | 0.4218 |
| Rack1 | 1.2E-194 | 0.3314 |
| Sh2d1a | 4.1E-136 | 0.3279 |
| Klra7 | 1.1E-32 | 0.3151 |
| Klf4 | 7.6E-97 | 0.3086 |
| Npm1 | 1.2E-75 | 0.2939 |
| Naca | 8.6E-107 | 0.2742 |
| Limd2 | 4.6E-80 | 0.2688 |
| Vim | 3.2E-37 | 0.2627 |
| Pycard | 1.5E-55 | 0.2487 |
| Hspe1 | 1.2E-55 | 0.2484 |
| Gas5 | 7.7E-71 | 0.2441 |
| Sell | 8.4E-63 | 0.2433 |
| Cmtm7 | 6.0E-66 | 0.2344 |
| Actb | 1.2E-40 | 0.2290 |
| Ms4a4b | 7.1E-63 | 0.2251 |
| Irf1 | 7.0E-34 | 0.2217 |
| Hsp90ab1 | 4.3E-53 | 0.2199 |
| Egr1 | 9.4E-34 | 0.2163 |
| Hspa8 | 1.8E-53 | 0.2157 |
| Il21r | 2.1E-53 | -0.2789 |
| Btg1 | 1.9E-76 | -0.2797 |
| Ubald2 | 1.2E-32 | -0.2893 |
| Hist1h1c | 2.2E-16 | -0.2966 |
| Srgn | 2.2E-99 | -0.2970 |
| Osgin1 | 6.4E-64 | -0.3035 |
| Klra9 | 5.7E-33 | -0.3312 |
| Cma1 | 8.5E-45 | -0.3348 |
| Dennd4a | 1.7E-104 | -0.3838 |
| S100a4 | 3.9E-40 | -0.3872 |
| Xcl1 | 4.8E-16 | -0.4006 |
| Serpinb9 | 7.3E-64 | -0.4076 |
| Bcl2a1b | 7.5E-70 | -0.4348 |
| Gzma | 7.3E-125 | -0.4483 |
| Prf1 | 4.5E-76 | -0.4488 |
| Neurl3 | 2.7E-111 | -0.4535 |
| Ccl3 | 9.9E-33 | -0.4703 |
| Ly6c2 | 1.2E-49 | -0.4725 |
| Dusp2 | 1.7E-122 | -0.5391 |
| Cd3g | 5.0E-23 | -0.6583 |
| Gzmc | 3.5E-20 | -0.7619 |
| Rgs1 | 1.9E-75 | -0.8273 |
| Ccl4 | 6.4E-136 | -0.8537 |
| Gzmb | 1.9E-266 | -0.9324 |

| cNK_5 | | |
|---|---|---|
| gene | p_val | avg_logFC |
| Klf2 | 0 | 1.1708 |
| Fos | 0 | 0.7854 |
| Irf1 | 0 | 0.6834 |
| Zeb2 | 0 | 0.6442 |
| Fosb | 0 | 0.6208 |
| Atf3 | 1.1E-156 | 0.5998 |
| Zfp36 | 0 | 0.5331 |
| Dusp1 | 5.8E-303 | 0.5207 |
| Sgk1 | 1.0E-129 | 0.4821 |
| Klf6 | 3.9E-292 | 0.4659 |
| Cma1 | 7.1E-286 | 0.4608 |
| Ly6c2 | 5.9E-123 | 0.4373 |
| Hist1h1c | 2.1E-94 | 0.4182 |
| Jun | 1.2E-233 | 0.4177 |
| Btg2 | 1.6E-257 | 0.4002 |
| Rhob | 8.1E-174 | 0.3897 |
| Ier2 | 1.1E-227 | 0.3756 |
| Egr1 | 3.1E-106 | 0.3690 |
| Icam1 | 2.2E-86 | 0.3601 |
| Ifng | 7.0E-67 | 0.3595 |
| Ccl3 | 2.6E-107 | 0.3517 |
| Rgcc | 6.4E-72 | 0.3459 |
| Vim | 1.6E-170 | 0.3383 |
| Gm26532 | 2.1E-147 | 0.3317 |
| Neat1 | 2.3E-102 | 0.3144 |
| Mpp7 | 4.9E-110 | -0.3370 |
| Smad7 | 6.6E-111 | -0.3463 |
| Tgfb1 | 4.1E-100 | -0.3497 |
| Osgin1 | 8.4E-140 | -0.3575 |
| Litaf | 1.9E-157 | -0.3614 |
| Gzmb | 1.2E-20 | -0.3652 |
| Hspa1b | 1.8E-17 | -0.3687 |
| Trbc1 | 1.9E-127 | -0.3707 |
| Emb | 1.1E-61 | -0.3729 |
| Ppp1r16b | 1.1E-164 | -0.3798 |
| Gm36723 | 4.0E-149 | -0.4366 |
| Prdx6 | 1.2E-180 | -0.4464 |
| Trbc2 | 6.0E-79 | -0.4477 |
| Bcl2a1b | 6.8E-117 | -0.4627 |
| Cd7 | 6.9E-146 | -0.4666 |
| S100a4 | 2.6E-86 | -0.5067 |
| Ly6e | 1.2E-191 | -0.5116 |
| Cd160 | 1.4E-164 | -0.6089 |
| Ltb | 1.2E-245 | -0.6932 |
| Rgs1 | 4.3E-58 | -0.7234 |
| Xcl1 | 8.4E-235 | -0.8050 |
| Hspa1a | 1.1E-13 | -0.8416 |
| Gzmc | 9.0E-66 | -0.8481 |
| Cd3g | 3.6E-124 | -0.9860 |

| cNK_6 | | |
|---|---|---|
| gene | p_val | avg_logFC |
| Ctla2a | 0 | 0.9929 |
| Emb | 0 | 0.8557 |
| Plac8 | 0 | 0.8522 |
| Gzmc | 1.8E-23 | 0.7968 |
| Tgfb1 | 0 | 0.5893 |
| Ccr2 | 0 | 0.5689 |
| Cotl1 | 0 | 0.5369 |
| Bst2 | 2.8E-218 | 0.5318 |
| Car2 | 0 | 0.5229 |
| Prdx6 | 0 | 0.5204 |
| Pmepa1 | 0 | 0.4944 |
| Vegfa | 0 | 0.4826 |
| Smad7 | 0 | 0.4807 |
| Socs3 | 0 | 0.4636 |
| Cxcr4 | 0 | 0.4593 |
| Runx3 | 6.6E-272 | 0.4538 |
| Cmtm7 | 0 | 0.4293 |
| Gramd3 | 0 | 0.4274 |
| Npm1 | 1.6E-251 | 0.4256 |
| Rack1 | 0 | 0.4255 |
| Ms4a4b | 3.7E-253 | 0.4230 |
| Stat3 | 7.6E-242 | 0.4084 |
| Shisa5 | 0 | 0.4059 |
| Gas5 | 2.7E-270 | 0.3943 |
| Aldoa | 1.0E-119 | 0.3850 |
| Emp3 | 1.2E-240 | -0.4158 |
| Sgk1 | 3.8E-81 | -0.4389 |
| Klra9 | 2.1E-117 | -0.4677 |
| Zeb2 | 3.9E-150 | -0.4781 |
| Dnaja1 | 6.4E-166 | -0.4800 |
| S100a6 | 6.2E-193 | -0.4903 |
| Ccl5 | 0 | -0.5124 |
| Gzma | 2.3E-290 | -0.5321 |
| S100a4 | 6.8E-137 | -0.5401 |
| Klf6 | 5.7E-251 | -0.5619 |
| Cma1 | 2.1E-241 | -0.5886 |
| Btg2 | 0 | -0.5891 |
| Dusp1 | 6.9E-284 | -0.6394 |
| Ly6c2 | 8.4E-183 | -0.6950 |
| Jun | 1.9E-195 | -0.7015 |
| Rgs1 | 4.6E-94 | -0.7928 |
| Cd3g | 7.8E-65 | -0.7928 |
| Dnajb1 | 1.2E-230 | -0.7980 |
| Ccl4 | 0 | -0.9348 |
| Fos | 2.0E-199 | -0.9525 |
| Klf2 | 4.2E-288 | -1.0461 |
| Ccl3 | 0 | -1.1192 |
| Hspa1a | 8.0E-208 | -1.4513 |
| Hspa1b | 0 | -1.6418 |

**Supplementary Table 1:** Top 25 upregulated genes and downregulated genes for each trILC1 and cNK cluster identified from scRNA. p-val = p value, avg_logFC = average log2(Fold Change).
